# Recurrent human papillomavirus-related head and neck cancer undergoes metabolic re-programming and is driven by oxidative phosphorylation

**DOI:** 10.1101/2020.08.21.261206

**Authors:** Avani Vyas, R. Alex Harbison, Daniel Faden, Mark Kubik, Drake Palmer, Qing Zhang, Hatice U. Osmanbeyoglu, Eduardo Méndez, Umamaheswar Duvvuri

## Abstract

Human papillomavirus (HPV) infection drives the development of some head and neck cancer squamous cell carcinomas (HNSC). This disease is rapidly increasing in incidence worldwide. Although these tumors are sensitive to treatment, ~10% of patients fail therapy. However, the mechanisms that underlie treatment failure remain unclear. Here, we show that the oxidative phosphorylation (OXPHOS) pathway is enriched in recurrent HPV-associated HNSC and may contribute to treatment failure. Nrf2-enriched HNSC samples from the Cancer Genome Atlas with enrichment in OXPHOS, fatty acid metabolism, Myc, Mtor, ROS, and glycolytic signaling networks exhibited worse survival. HPV-positive HNSC cells demonstrated sensitivity to the OXPHOS inhibitor, IACS-010759, in a Nrf2-dependent manner. Further, using murine xenograft models, we identified Nrf2 as a driver of tumor growth. Mechanistically, Nrf2 drives ROS and mitochondrial respiration, and Nrf2 is a critical regulator of redox homeostasis that can be crippled by disruption of OXPHOS. Nrf2 also mediated cisplatin sensitivity in endogenously overexpressing primary HPV-related HNSC cells. Cisplatin treatment demonstrated Nrf2-dependent synergy with OXPHOS inhibition. These results unveil a paradigm shifting translational target harnessing Nrf2-mediated metabolic reprogramming in HPV-related HNSC.

## Introduction

While human papillomavirus (HPV)-related head and neck squamous cell carcinoma (HNSC) responds favorably to concurrent platinum-based chemoradiation (1), treatment failure portends a grim prognosis with limited treatment options including morbid surgical resection, chemotherapy (i.e., with platinum-based agents) and/or reirradiation, or clinical trials mainly with immunotherapeutic agents (2). From a public health standpoint, the incidence of HPV-related HNSC has surpassed that of cervical cancer and is projected to continue rising until at least 2060 (3,4). Given the ongoing rise in HPV-related HNSC and challenges of managing treatment non-responders, it is critical to understand the biological mechanisms underlying recurrent HPV-related HNSC so that we may mitigate recurrence at the time of initial therapy.

To better understand the signaling the mechanisms that drive recurrent HPV-related HNSC, our group sought to identify differences in the genomic landscapes between metachronous recurrent and primary HPV-related HNSCs (5). We had previously shown that metachronous recurrent HPV-related HNSCs shared a genomic landscape with aggressive smoking- and alcohol-associated (HPV-unrelated) HNSCs. A subsequent study using gene expression data, identified a subset of primary HPV-related HNSCs that exhibited a poor treatment response and shared molecular similarities with HPV-unrelated HNSCs (6). In comparison, findings from our prior genomic analysis revealed gene mutations exclusive to primary HPV-related HNSCs that recurred relative to primary HPV-related HNSC cases that did not recur (i.e., *NFE2L2, TSC2, BRIP1*, and *NBN*). Genomic alterations of HPV-unrelated HNSCs, as characterized by the Cancer Genome Atlas (TCGA) study include loss-of-function *TP53* mutations and *CDKN2A* inactivation in addition to a key, and underappreciated, role of the transcription factor *NFE2L2* (the gene which encodes Nrf2) (7). *NFE2L2* plays a role in regulating oxidative stress and mitigating the efficacy of chemoradiation (8) and interacts with the HPV E1 protein (9).

While prior work has focused on uncovering the molecular characteristics of primary HPV-related HNSC that portend a poor prognosis, we continue to experience a paucity of treatment options in the management of recurrent disease that tend to have high morbidity. In this study, we sought to elucidate targetable biological mechanisms underlying recurrent HPV-related HNSC. Our overarching hypothesis was that HPV-related HNSCs would demonstrate differential expression in key signaling pathways which may be exploited as therapeutic targets. Intriguingly, we observed activation of the OXPHOS pathway in the background of metabolic gene dysregulation among matched patient samples from primary and subsequent metachronous recurrent tumors. We further confirmed that Nrf2 functions as a driver of growth in an OXPHOS-dependent manner in HPV-associated HNSC. *In vitro* findings illustrated a functional dependence on OXPHOS and ROS among Nrf2-overexpressing cells conferring a critical weakness to OXPHOS inhibition. Taken, together, these findings implicate Nrf2 and OXPHOS as drivers of a subset of HPV-associated HNSC and raise the intriguing possibility that Nrf2 may serve as a novel therapeutic target in this subset of patients.

## Results

### Patient Characteristics

Starting with ten patients with matched primary (pOPSCC) and metachronous recurrent HPV-related oropharyngeal squamous cell carcinoma (rOPSCC) from the University of Pittsburgh, our first aim was to identify transcriptional differences between the pOPSCC and rOPSCC to gain mechanistic insight into the evolutionary adaptations of metachronous recurrent tumors (**Figure 1A**). The median age of the cohort was 60 years old (**Supplemental Table 1**). Based on the AJCC 8^th^ edition staging manual (10), seven of ten patients presented with stage I disease and three out of ten with stage II disease. Of the 10 patients in the cohort, six were never smokers, three patients had a greater than 10 years smoking history, and one patient had a 15-year history of chewing tobacco use. Median overall survival was 36.4 months among the cohort.

**Figure 1.**
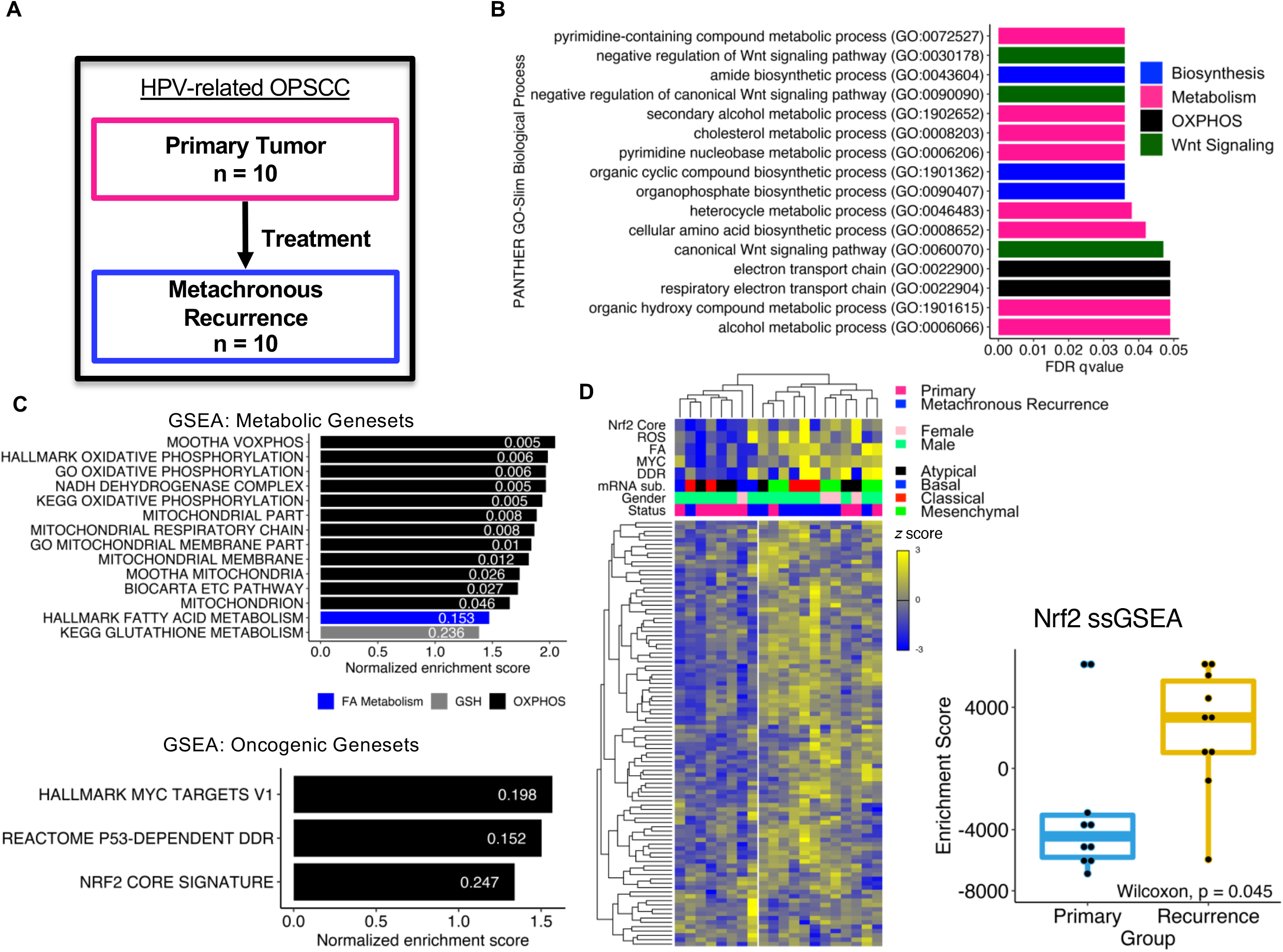
Nrf2 and OXPHOS signaling pathways are enriched in metachronous recurrent (rOPSCC) HPV-related oropharyngeal squamous cell carcinoma. (**A**) RNA expression assays performed on ten paired primary OPSCC (pOPSCC) and rOPSCC. (**B**) PANTHER pathway analysis of differentially expressed genes comparing rOPSCC versus pOPSCC. Pathways with over-representation test false discovery rate (FDR) q-value < 0.05 are displayed. (**C**) Gene set enrichment analysis (GSEA) of rOPSCC versus pOPSCC. Normalized enrichment scores (NES) plotted. FDR q-values labeled within the bars. *Upper plot:* GSEA using panel of metabolic gene sets. *Lower plot:* GSEA using panel of oncogenic gene sets. (**D**) *Left panel:* Hierarchical clustering of hallmark OXPHOS gene set among pOPSCC and rOPSCC. Rows represent genes. Columns represent tumors. Row and column dendrograms demonstrate clustering results. Normalized expression counts in cells. Color bars in the upper plot illustrate single sample GSEA (ssGSEA) enrichment score. *mRNA sub*., head and neck cancer molecular subtype classification; *ROS*, hallmark reactive oxygen species; *FA*, hallmark fatty acid metabolism; *MYC*, hallmark *MYC* v1; *DDR*, Reactome p53-dependent DNA damage repair; *Nrf2 Core*, Nrf2 core signature (ref. 11). *Right panel:* Nrf2 ssGSEA enrichment scores between groups.

### Nrf2 and OXPHOS signaling pathways are enriched in metachronous recurrent HPV-related OPSCCs

PANTHER pathway analysis revealed a preponderance of biological processes involved with biosynthesis, metabolism, oxidative phosphorylation (OXPHOS), and Wnt signaling (FDR q < 0.05; **Figure 1B**). To further understand the mechanisms driving recurrent OPSCC, we performed gene set enrichment analysis (GSEA) utilizing pre-specified metabolic and oncogenic gene sets (**Supplemental Tables S2 and S3**) based on our prior research (5) and *a priori* knowledge of HPV-unrelated HNSC biology. This demonstrated selective enrichment of OXPHOS pathways among the rOPSCCs for the metabolic gene sets (FDR q < 0.25; **Figure 1C**, *upper panel*). In the oncogenic GSEA, rOPSCCs were enriched in Myc, p53-dependent DNA damage repair, and Nrf2 core signaling (11) pathways (FDR q < 0.25; **Figure 1C**, *lower panel*). Myc plays a critical role in regulating both glycolysis and OXPHOS. Nrf2 canonically regulates the antioxidant response element (ARE) genes in response to oxidative stress but also plays a critical role in OXPHOS. We next performed hierarchical clustering of expression data restricted to genes from the hallmark OXPHOS gene set revealing two main clusters (**Figure 1D**, *left panel*). One cluster included eight of the ten rOPSCCs while the second cluster was populated by six of the ten pOPSCCs. To assess for genomic correlations with OXPHOS gene expression in these data, we performed single sample GSEA (ssGSEA) on the paired primary and recurrent tumor samples using oncogenic gene sets that were significantly enriched among the rOPSCCs (**Figure 1D**, bars above main heatmap). We also performed ssGSEA using the Hallmark fatty acid metabolism gene set which was enriched in the metabolic GSEA and Hallmark reactive oxygen species hypothesizing that these genes would be associated with Nrf2 target gene expression. We compared the enrichment scores between the primary and recurrent tumors for each gene set and observed significantly greater enrichment scores among the rOPSCCs for the Nrf2, fatty acid metabolism, and Myc gene sets (Wilcoxon rank-sum p < 0.05; **Figure 1D**, *right panel*; **Supplemental Figure 1**). In addition, we performed a molecular subtype analysis to assess correlation between the primary and recurrent OPSCCs with previously defined centroids (12) hypothesizing that the rOPSCCs would share characteristics with the classical (i.e., smoking-related) subtype. We found that eight of the ten rOPSCCs clustered with classical (4/10) or mesenchymal (4/10) mRNA subtypes while the majority of pOPSCCs clustered with the atypical (5/10) or basal subtypes (2/10; **Figure 1D**, *left panel*).

To further assess context-specific gene regulatory networks that may be enriched or diminutive in rOPSCCs, we inferred transcription factor (TF) activities based on RNA-seq data (13). We observed a significant increase in *E2F3* and *SMAD3* activity among the rOPSCCs (Wilcoxon rank-sum, p < 0.05; **Supplemental Figure 2**). *BACH1/NFE2/NFE2L2* activity was increased in the rOPSCCs though not statistically significantly (Wilcoxon rank-sum, p = 0.062). *PKNOX1/TGIF2* activity was significantly decreased among the rOPSCCs (Wilcoxon rank-sum, p = 0.006). Interestingly, Nrf2 activation contributes to epithelial-mesenchymal transition (EMT) by mediating a decrease in E-cadherin and stimulating TGF-β1-induced *SMAD2/3* activity (14). In turn, TGF-β signaling is involved in stimulating both glycolysis and mitochondrial respiration. As *TGIF2* represses transcription of TGF-β-responsive genes by recruiting histone deacetylases (15), it follows that the *TGIF2* signaling network was decreased in the rOPSCCs while *SMAD3* activity was increased.

### Nrf2 activation portends worse survival among HPV-related HNSCs

To investigate the effect of Nrf2 in a separate cohort, we utilized the TCGA head and neck cancer data (7). With a set of 99 HPV-positive TCGA HNSC samples, we performed a survival analysis stratified by Nrf2 or Myc genomic alteration status defined as expression > 2, mutated, or copy number gain or amplification. Mutation status was included given that Nrf2 and Myc somatic variants in the TCGA HNSC data are either characterized oncogenic hotspot mutations or predicted to have a functional impact. Surprisingly, we identified a significant difference in survival among the Nrf2-altered (*Nrf2 Up*) samples (log-rank test, p = 0.005) versus no difference in survival among the Myc-altered (*Myc Up*) samples (log-rank test, p = 0.16; **Figure 2A**). These data suggest that Nrf2, but not Myc, may functionally drive OXPHOS in HPV-associated HNSC. Next, we sought to identify if there were signaling network-level relationships with survival among the TCGA HPV-related HNSC samples.

**Figure 2.**
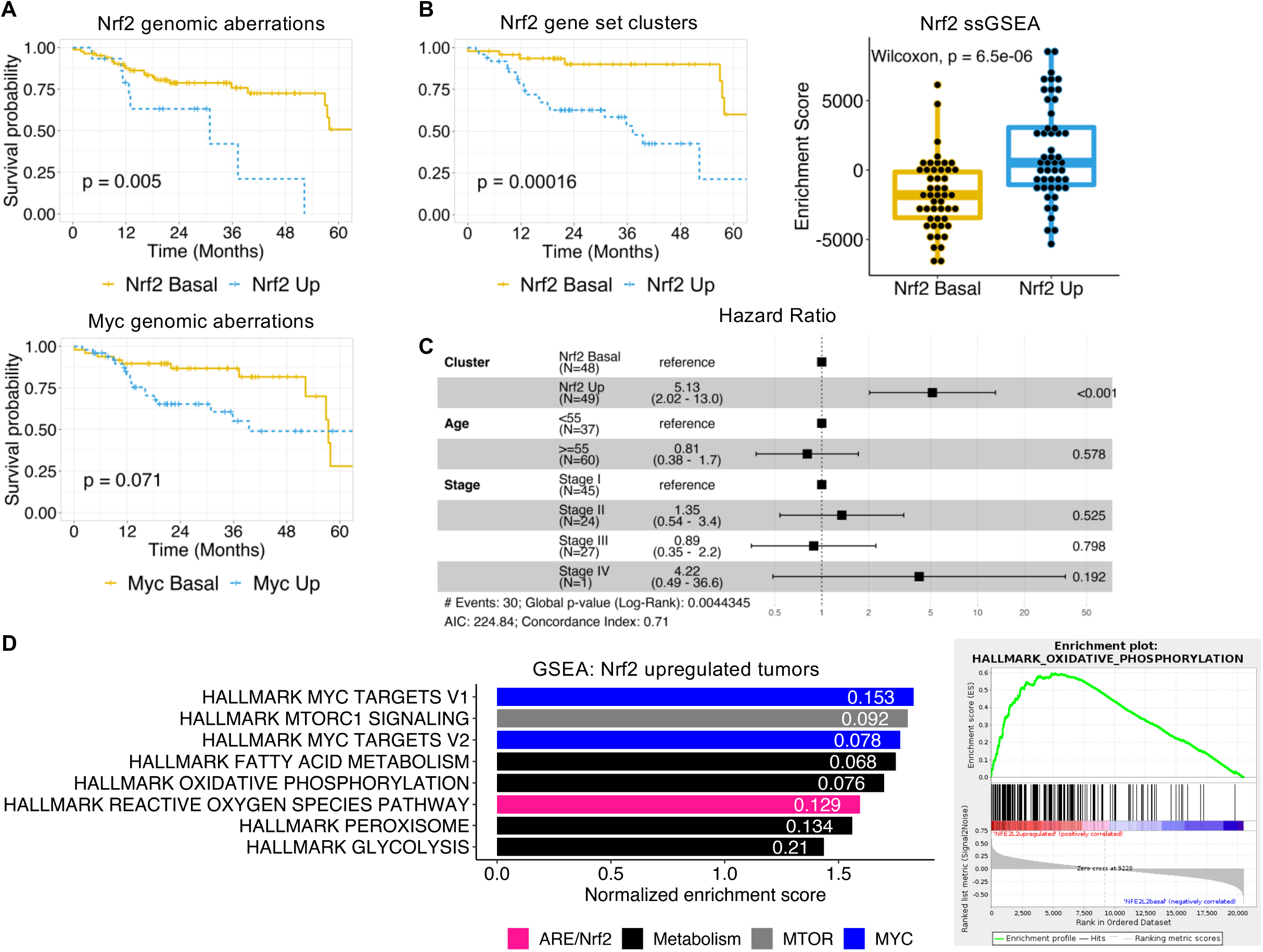
Nrf2 activation decreases survival among TCGA HPV-related head and neck squamous cell carcinomas (HNSC). (**A**) Survival stratified by Nrf2 (*top panel*) or Myc (*bottom panel*) genomic alteration status (*Up*: gene expression > 2, mutation, copy number gain or amplification; *Basal*: gene expression ≤ 2, no mutation, copy neutral). Log-rank global p-value labeled. (**B**) *Left panel*: Survival stratified by Nrf2 status as determined by clustering TCGA HPV-related HNSC samples on Singh *NFE2L2* gene set expression. Log-rank p-value labeled. *Right panel*: Singh *NFE2L2* ssGSEA enrichment scores stratified by gene set cluster membership as assigned in clustering analysis. (**C**) Cox regression hazard ratio by Nrf2 gene set expression cluster, age group, and tumor stage (AJCC 8th edition, clinical staging). Point estimates plotted with 95% confidence intervals represented by whiskers. P-values for each hazard ratio by predictor are represented on the far right of plot. (**D**) *Left panel*: GSEA of TCGA HPV-related HNSC Nrf2 upregulated tumors as defined in (**A**; *top panel*). FDR q-values labeled within bars. *Right panel:* Hallmark oxidative phosphorylation gene set enrichment plot.

We performed clustering analyses on the TCGA expression data selecting genes within gene sets of interest using the Consensus Cluster Plus algorithm. Cluster assignments were used for stratified survival analyses. Then, single sample GSEA was performed to quantify enrichment scores of tumor samples allowing us to infer the correlation between a respective cluster gene set enrichment and survival (e.g., high enrichment with worse survival). Using the Singh *NFE2L2* targets gene set (16) for clustering the TCGA HPV-related HNSC expression data, we found a significant difference in survival with the Nrf2-enriched cluster (*Nrf2 Up*) conferring worse survival (log-rank test, p = 0.00016; **Figure 2B**). When controlling for Nrf2 enrichment cluster, age, and stage, the Nrf2-enriched status conferred a 5.13-fold (Hazard ratio 95% CI: 2.02 – 13.0; p < 0.001) increased risk of mortality (**Figure 2C**). GSEA using the Hallmark gene sets was performed comparing the Nrf2-altered vs -unaltered TCGA HPV-related HNSC tumors as defined by genomic alteration status in **Figure 2A**. Interestingly, Nrf2-altered tumors featured enrichment in *MYC, MTOR*, fatty acid metabolism, OXPHOS, ROS, peroxisome, and glycolysis gene sets suggesting a relationship between Nrf2 and critical metabolic signaling pathways (**Figure 2D**).

While Nrf2 appeared to play an integral role in the metachronous recurrent setting, we were interested in testing for an independent or synergistic effect of Nrf2 transcriptional targets and other critical metabolic regulatory genes on survival. We performed a survival analysis among the TCGA HPV-positive HNSC tumors stratified by Nrf2 genomic alteration status in combination with Hmox1, Nqo1, or Sqstm1 genomic alterations (defined by gene expression > 2, copy number gain or amplification). Interestingly, combined Nrf2- and Hmox1-altered tumors had the worst survival among the Nrf2 ± Hmox1 alteration group while Nrf2-altered in combination with Nqo1- or Sqstm1-altered tumors did not represent the worst survival in their respective strata (**Supplemental Figure 3**). We also tested if Hmox1, Nqo1, or Sqstm1 upregulation had an independent association with survival among the TCGA HPV-related HNSCs. Only tumors with upregulation of Hmox1 had statistically significantly worse survival compared to those without Hmox1 upregulation (**Supplemental Figure 4**). These data implicate OXPHOS as a metabolic driver of HPV-related HNSC and suggest that recurrent/metastatic tumors co-opt this pathway during disease progression.

We further tested this hypothesis by evaluating the effect on survival based on metabolic and oncogenic signaling network genomic alterations among the TCGA HPV-related HNSC samples. We performed clustering on the expression data followed by survival analyses stratified by cluster. Single sample GSEA was used to assess cluster enrichment as described above for gene sets including OXPHOS, ROS, fatty acid metabolism, DNA damage response, and Myc. These gene sets were included as they were significantly enriched in the Pittsburgh recurrent tumors. The ROS gene set was tested as it includes genes involved in oxidative stress response. Interestingly, clusters with the greatest enrichment did not confer the worst survival among the OXPHOS and ROS gene sets (**Supplemental Figure 5 and 6**). Similarly, clusters with the highest enrichment among the fatty acid metabolism, DNA damage response, and Myc gene sets did not exhibit the worst survival (**Supplemental Figure 7 and 8**). We repeated the above survival analyses in the TCGA HPV-unrelated (i.e., smoking-related) HNSC samples. There was not a significant difference in survival when stratified by Nrf2- or Myc-altered or gene set cluster status (data not shown). However, there was a significant difference in survival when stratified by OXPHOS cluster status (data not shown). Taken together, our genomic analyses identify and association between Nrf2 and metabolic reprogramming with the development and progression of HPV-related HNSC. Thus, we sought to characterize the mechanisms by which Nrf2 drives tumorigenesis in HPV-related head and neck cancer.

### Nrf2 promotes cell proliferation in HPV-positive HNSC cell lines

Next, we sought to investigate the effect of Nrf2 overexpression on growth in HPV-related HNSC cells lines given findings in our rOPSCC and the TCGA HPV-related HNSC data which suggested that Nrf2 is involved with recurrence and worse overall outcomes, respectively. In order to establish a stable Nrf2 overexpressing cell line, HPV-16-positive UMSCC47 and UPCI-SCC90 were infected with a Nrf2-expressing retrovirus (murine stem cell virus [MSCV]-Nrf2). These cell lines were chosen given their minimal endogenous Nrf2 expression (**Supplemental Figure 9A**). The efficacy of Nrf2 overexpression was assessed by performing qRT-PCR (**Figure 3A**) on exponentially growing cells in different passages. The proliferative potential of Nrf2 upregulation in these cells was measured by WST-1 (data not shown) and a colony formation assay (CFA; **Figure 3B**). To test the effect of Nrf2 knockdown on cell growth, we utilized an HPV-16-positive cell line with increased endogenous expression of Nrf2 (93VU147T). We transiently knocked down Nrf2 using short interfering RNA (siRNA) and measured the growth index using a CFA (area of CFA for siNrf2 relative to control: ~50%, p = 0.01, n = 2, **Figure 3C**). To assess the effect of an exogenous Nrf2 inhibitor, we treated 93VU147T cells with the coffee alkaloid, trigonelline (trig), observing attenuated proliferation (67% proliferation with trig relative to control, n = 3, p < 0.0001; data not shown).

**Figure 3.**
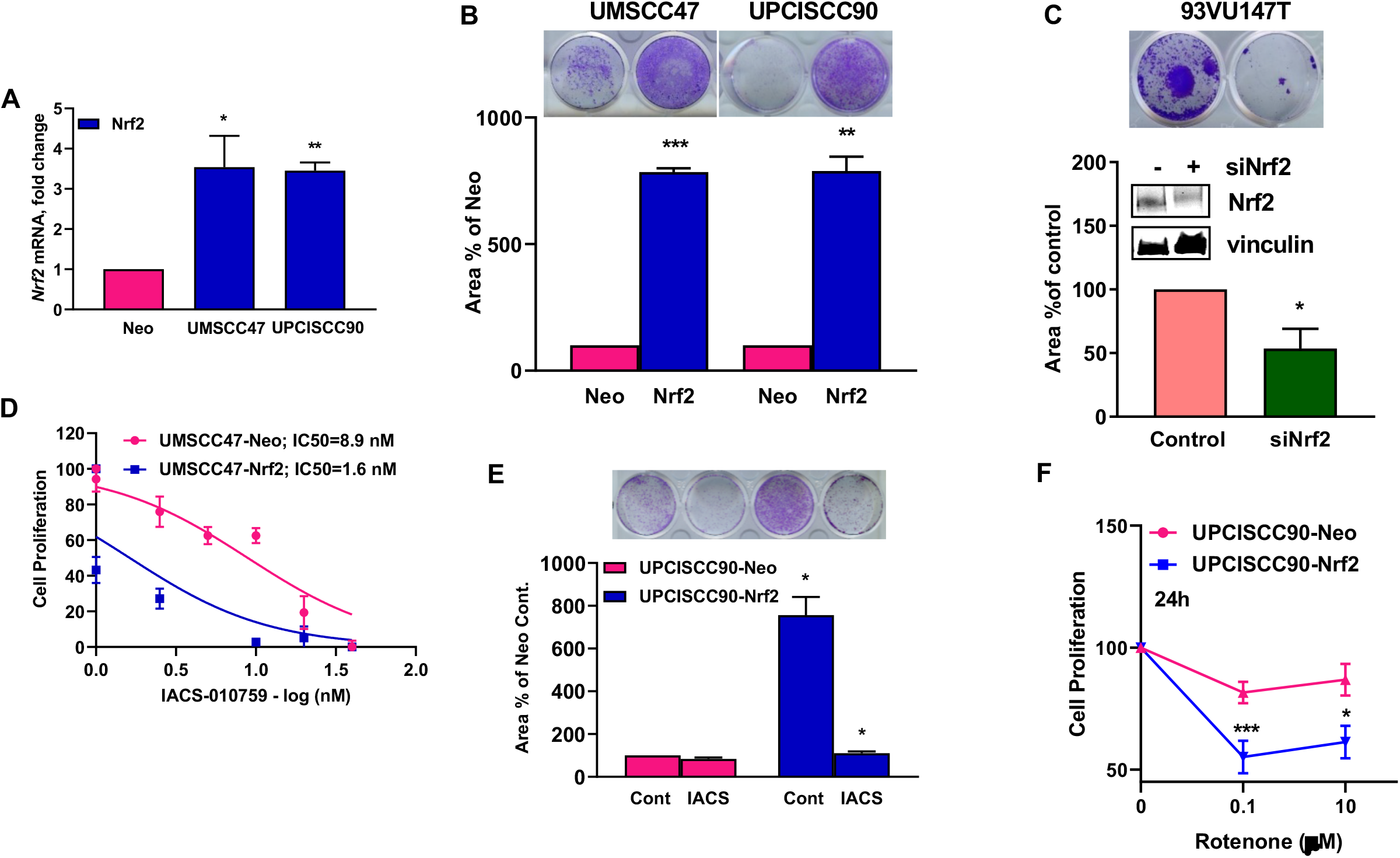
Nrf2 promotes cell proliferation in OXPHOS-dependent manner in HPV-positive HNSC cells. (**A**) qPCR demonstrates overexpression of Nrf2 after retroviral stable infection. (**B**) Representative image (upper) of colony formation assay (CFA) and quantification (lower) in stably expressing empty vector (Neo) or Nrf2 cells. (**C**) Representative picture (upper) of CFA and quantification (lower) in 93VU147T after transient Nrf2 knockdown. (**D**) Cell proliferation assay for IACS-010759 (IACS)-treated UMSCC47-Neo and -Nrf2 cells measured after 24h of treatment (representative data from 3 experiments). (**E**) CFA after treatment with IACS in UPCISCC90-Neo and - Nrf2 cells. (**F**) Cell proliferation in UPCISCC90-Neo and Nrf2-cells after treatment with 0.1 or 10 μM rotenone. Results represent mean ± standard error of the mean (SEM).

Nrf2 has recently emerged as a key regulator of mitochondrial function. Mitochondria generate ATP through OXPHOS which is the primary energy source of cells in their basal state. In contrast, dividing cells including tumors and activated lymphocytes utilize aerobic glycolysis to optimize substrate production for generating new cells. However, our transcriptional analyses suggested a predilection for OXPHOS gene expression in tumors with poor survival characteristics. Therefore, we tested IACS-01759 (IACS), a potent and selective electron transport chain complex I inhibitor which is currently in phase I trials. IACS in nM concentrations was significantly more cytotoxic to UMSCC47-Nrf2 cells (IC50=1.6nM) compared to Neo cells (IC50=8.9 nM, n=3, **Figure 3D**).

We confirmed the inhibition of cell proliferation by IACS in another cell line harboring stably expressed Nrf2 in UPCISCC90 cells (**Figure 3E**; **Supplemental Figure 9A** and **B**). To further evaluate the role of OXPHOS in cellular proliferation, we tested the effect of rotenone via CFA in UPCISCC90-Neo and -Nrf2 cells (data not shown). Rescuing the cells from rotenone after 2h treatment was irreversible in the Nrf2-dependent cells and the cells failed to form colonies even after 10 days in culture.

The mitochondrial toxin, rotenone, also inhibited cellular proliferation in a Nrf2-dependent manner within 2h treatment at a low dose of 0.1μM (data not shown). This effect was more pronounced in the Nrf2-overexpressing cells, as observed at 24h (~30% reduction, ***p<0.0001, n=3; **Figure 3F**). A comparison of IC50 values in different Nrf2-overexpressing cell lines (**Supplemental Figure 9C**) suggests that Nrf2 upregulated cells are heavily reliant on OXPHOS for growth and survival in contrast to Nrf2 basal cells. Taken together, these data infer that Nrf2 promotes cell proliferation in an OXPHOS-dependent manner in HPV-positive HNSC.

### Nrf2 overexpressing xenografts exhibit enhanced growth

The *in vivo* effect of UMSCC47 stably expressing Nrf2 and Neo was assessed in a NOD SCID mouse model. Cells were injected into both flanks and allowed to establish for a week. The mean tumor weight and volume increased significantly over time in the mice implanted with Nrf2 cells relative to Neo. At day 21 post-implantation, the tumor length and width had reached more than 20 mm in the Nrf2 xenografts. The Nrf2 expressing tumors had 3-fold greater tumor weight at time of harvest (p < 0.05, n = 3 or 4, **Figure 4A** and **4B**). The tumor volume of Nrf2 overexpressing cells amplified by 3.5-fold at day 7 (p < 0.05), 6-fold at day 14, and 14-fold at day 21 (p<0.0001; **Figure 4C**). These data advocate that Nrf2 expressing tumors have enhanced proliferative advantage over the low-Nrf2 expressing tumors.

**Figure 4.**
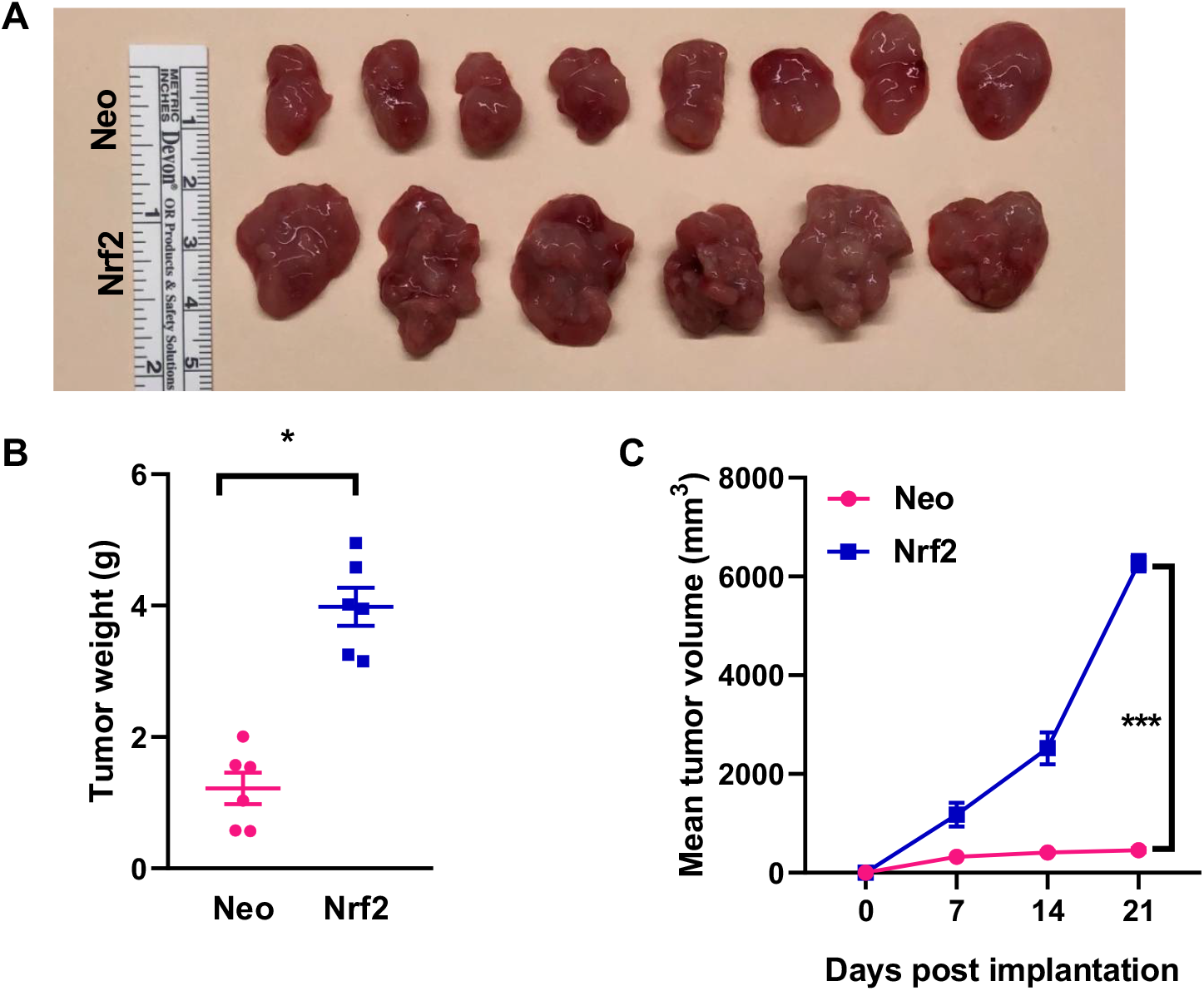
Nrf2 overexpressing xenografts exhibit enhanced growth. Both flanks of NOD/SCID mice were injected subcutaneously with UMSCC47-Neo and -Nrf2 cells and tumor volumes were measured weekly. (**A**) Images of harvested UMSCC47 tumors are demonstrated for each group. (**B**) Tumor weights collected on day 21 post-implantation. Difference in mean tumor weights between groups compared using Student’s t-test. (**C**) UMSCC47 tumor growth over time.

### Nrf2 activation upregulates mitochondrial respiration in a ROS-dependent manner

In order to gain insight into the redox balance modulated by Nrf2 overexpression among HPV-positive HNSC cell lines, we measured markers of oxidative stress among control and Nrf2 overexpressing conditions hypothesizing that Nrf2 overexpression would abrogate oxidative stress. Given that ROS are a natural byproduct of mitochondrial respiration and that OXPHOS was increased in Nrf2-overexpressing cells in our prior experiments, we sought to investigate the impact of ROS on mitochondrial respiration. Nrf2-overexpressing cells demonstrated increased expression of the cytoprotective and detoxifying genes, heme oxygenase 1 (*HMOX1*) and NAD(P)H:quinone oxidoreductase-1 (*NQO1*) in UMSCC47 (3- and 5-fold, respectively; p-value < 0.05, n=3; **Figure 5A**) and UPCISCC90 cells (25- and 40-fold respectively; **Supplemental Figure 10A**). The amplification of *HMOX1* and *NQO1* were also confirmed in a Nrf2-enriched primary HPV-positive HNSC cell line (UPCI-UDSCC17-70; **Supplemental Figure 9D**).

**Figure 5.**
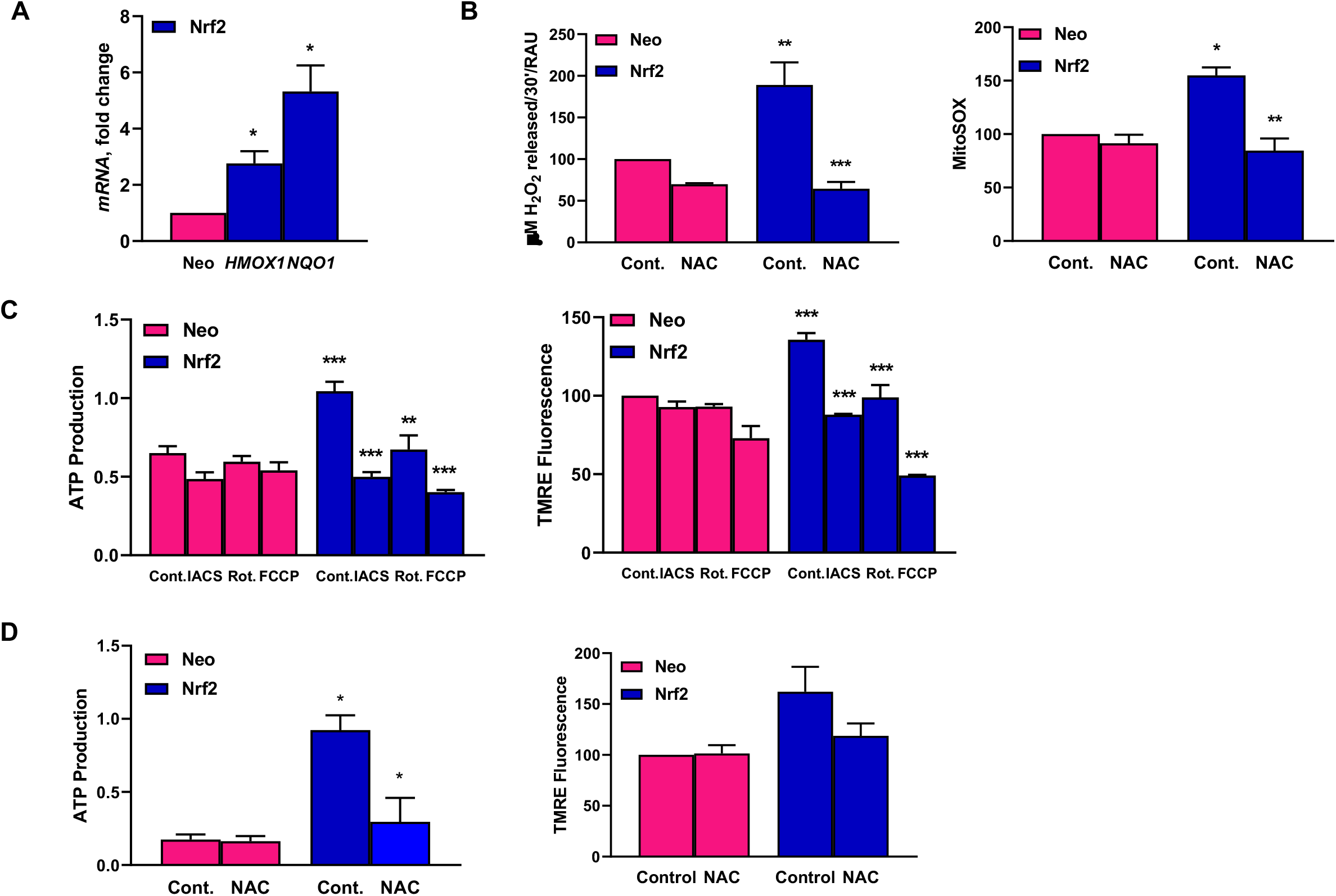
Nrf2 activation upregulates mitochondrial activity in a ROS-dependent manner. (**A**) qPCR for *HMOX1* and *NQO1* in Neo and Nrf2-expressing UMSCC47 cells. (**B**) *Left panel*: ROS (H2O2 released) measured in cells treated with and without 10mM N-acetyl cysteine (NAC) for 3h. *Right panel*: Mitochondrial superoxide measured in control versus 10mM NAC (3h)-treated cells via MitoSOX Red. (**C**) Nrf2 promotes ATP production and is dependent on mitochondria. *Left panel*: Nrf2 driven ATP production is abrogated upon treatment with mitochondrial inhibitors. *Right panel*: Mitochondrial membrane potential measured using TMRE dye. (**D**) Nrf2 drives ATP production via ROS production (*left panel*) and TMRE fluorescence (*right panel*) measured in cells treated with and without 10mM NAC (3h).

To quantify relative ROS production in Nrf2-enriched cells, we measured hydrogen peroxide (H_2_O_2_) and superoxide (O2^•−^) formation in intact and live cells using the Amplex Red and MitoSOX assays, respectively. We used the free radical scavenger *N*-acetylcysteine (NAC) to test the effect of abrogating ROS in Nrf2-enriched and Nrf2-basal cells. The basal rate of ROS production in UMSCC47-Nrf2 was almost double compared to parental cells as measured by the Amplex Red assay (p-value < 0.001, n=3; **Figure 5B**, *left panel*). NAC decreased ROS production to a greater degree in the Nrf2-overexpressing UMSCC47 cells (by ~60 and 70% respectively, p-value < 0.0001, n=3) relative to Neo diminishing ROS to basal levels. NAC did not significantly decrease ROS in cells expressing empty vector (**Figure 5**, *left panel*).

Next, we used MitoSOX Red in live cells to detect mitochondrial superoxide formation as an additional tool to quantify ROS production as a function of Nrf2 expression status. The results from combined data (n=3) indicate 50% more MitoSOX in Nrf2-enriched UMSCC47 cells compared to control cells (p-value < 0.05, n=3; **Figure 5B**, *right panel*). There was a significant reduction in MitoSOX levels in the Nrf2-overexpressing cells treated with NAC (~50%), but no significant changes between NAC versus control in Neo cells (**Figure 5B**, *right panel*). The Nrf2 inhibitor, trigonelline, decreased ROS and mitochondrial function in 93VU147T and UMSCC47-Neo and Nrf2 cells (data not shown) corroborating the role of Nrf2-mediated OXPHOS reprogramming and ROS signaling.

To further understand the sequelae of Nrf2-mediated OXPHOS reprogramming, we assessed the effect of Nrf2 overexpression on mitochondrial function by measuring ATP production and mitochondrial membrane potential. Compared to Neo cells, the Nrf2-overexpressing UMSCC47 cells exhibited ~1.7-fold higher ATP production which was mitigated after treatment with rotenone (p<0.001), IACS (~65% decrease, p < 0.0001), and FCCP (~40% decrease, p < 0.0001, n=3; **Figure 5C**, *left panel*). Next, we measured mitochondrial membrane potential in live cells using cell permeable tetramethylrhodamine ethyl ester (TMRE). The mitochondrial membrane potential is generated by proton pumps and is an essential component of energy storage during OXPHOS. The control UMSCC47-Nrf2 cells had greater TMRE fluorescence than control Neo cells (by ~35%, p < 0.0001), but this difference was mitigated significantly by the use of IACS or rotenone in the Nrf2-enriched cells (~50%, p < 0.0001, n=3; **Figure 5C**, *right panel*). Similarly, FCCP significantly decreased the mitochondrial membrane potential by 75% in the Nrf2 cells (p < 0.0001; **Figure 5C**, *right panel*). In contrast, ATP production and mitochondrial membrane potential in the Neo cells treated with IACS, rotenone, or FCCP were not significantly different compared to the control Neo cells.

To elucidate whether mitochondrial respiration in Nrf2 cells is dependent on ROS, we tested the effect of a free radical scavenger, NAC, on ATP production and mitochondrial membrane potential. NAC abrogated 70% (p < 0.0001, n=3; **Figure 5D**, *left panel*) of ATP production in Nrf2 cells, diminishing ATP production to that observed in Neo cells. In contrast, ATP levels were unchanged in Neo cells after NAC exposure (**Figure 5D**, *left panel*). Mitochondrial membrane potential did not change significantly after NAC treatment in Nrf2 cells exhibiting a ~30% decrease relative to control Nrf2 cells (**Figure 5D**, *right panel*). To confirm these findings were not exclusive to the UMSCC47 cell line, we repeated these experiments in the UPCISCC90 cell line stably overexpressing Nrf2 or empty vector. The results were consistent with the above findings (**Supplemental Figure 10**). Overall, these data suggest that increased ROS in the context of Nrf2 overexpression is not only a collateral phenomenon but may represent a positive feedback mechanism promoting OXPHOS.

### Nrf2-overexpression confers cisplatin sensitivity synergistically with OXPHOS inhibition in HPV-positive HNSC cells

Prior studies have demonstrated that Nrf2 can mediate resistance to chemotherapeutic agents including cisplatin (17-20). However, to the best of our knowledge, the role of NRF2 in HPV-related HNSC has not been studied. Since cisplatin is commonly used in the definitive or adjuvant setting in the treatment of HPV-related HNSC, we sought to investigate the effect of Nrf2 on cisplatin sensitivity in our models. Thus, we investigated the dose-dependency of cisplatin on cell proliferation comparing cells with endogenously elevated Nrf2 to those with genetically knocked-down Nrf2. The data shown in Figure 6A (n = 3) indicate that Nrf2 repression yields a ~7-fold lower IC50 of cisplatin in 93VU147T cells. Similar results were obtained using a primary cancer cell line (UPCI:UDSCC17-70) harboring endogenously elevated Nrf2 expression with or without siNrf2 (Figure 6B). Nrf2 knockdown was confirmed by qRT-PCR (Figure 6B, *left panel*). In siNrf2 cells treated with cisplatin, a 2.2-fold decrease in the IC50 was observed compared to control (n = 2, Figure 6B, *right panel*). As cisplatin induces oxidative stress, we hypothesized that co-treatment with IACS would be synergistically cytotoxic. To test this hypothesis, we treated UMSCC47-Neo and Nrf2 cells with cisplatin alone (CDDP) or in combination with IACS (Figure 6C). The combination synergistically sensitized the Nrf2 cells but not the Neo cells. Synergistic inhibition of growth was also observed with the combination of a Nrf2 inhibitor, trigonelline, and cisplatin in 93VU147T cells (data not shown). However, trigonelline was not as potent as IACS. These data indicate that Nrf2-overexpression mediates a delicate balance between ROS and antioxidant production through OXPHOS reprogramming suggesting a targetable weakness in Nrf2-enriched head and neck cancer.

**Figure 6.**
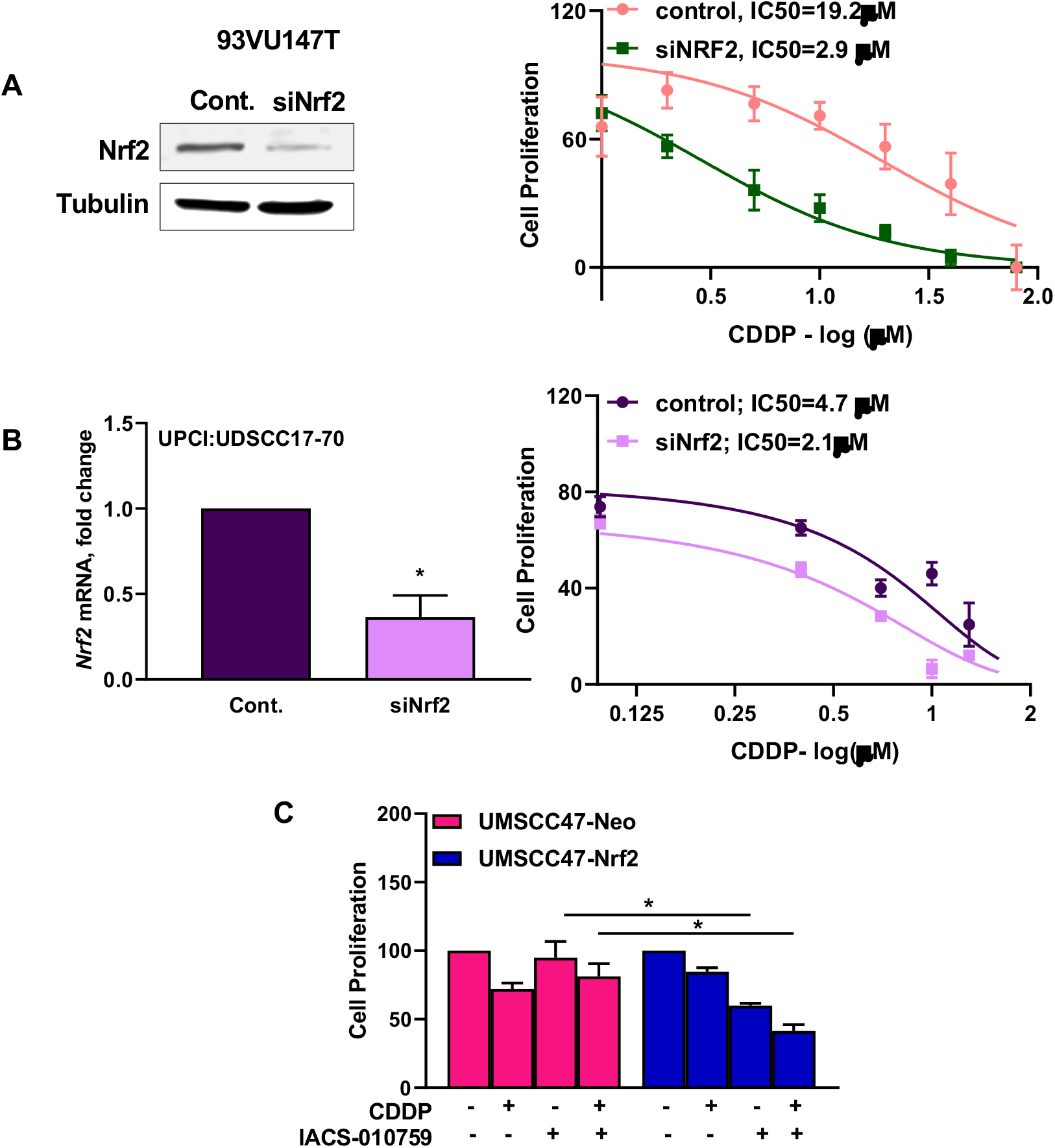
Nrf2 status mediates cisplatin sensitivity synergistically with OXPHOS inhibition in HPV-positive HNSC cells. (**A**) Western blot for Nrf2 in 93VU147T after transient transfection with siNrf2 for 24h (left panel). Cell proliferation in 93VU147T cells measured after Nrf2 knockdown (right panel). (**B**) Nrf2 knock-down sensitizes primary tumor cells to cisplatin. *Left panel*: qRT-PCR for Nrf2 in the primary cell line, UPCI:UDSCC17-70, after transient transfection with siNrf2 for 24h (left panel). *Right panel*: Cell proliferation in CDDP-treated UPCI:UDSCC17-70 cells in control and Nrf2 knockdown conditions. (C) Complex I inhibition is synthetically lethal with cisplatin in Nrf2 overexpressing cells. UMSCC47-Neo and -Nrf2 cells were treated with 10μM CDDP, 100 nM IACS, or combination for 24h. Differences in means calculated using one-way ANOVA with Tukey’s test.

## Discussion

The principal findings of this study were two-fold. First, gene expression analyses of paired metachronous recurrent versus primary HPV-related HNSCs demonstrated enrichment of the Nrf2 and OXPHOS signaling pathways amidst a milieu of metabolic gene dysregulation specific to the metachronous recurrences. Second, murine xenograft and *in vitro* analyses of HPV-related HNSC revealed Nrf2-dependent cellular proliferation. Further, we identified Nrf2-mediated OXPHOS dependency and sensitization to mitochondrial complex I inhibition with IACS-010759. Lastly, we found translational potential in Nrf2-mediated synergy between cisplatin and OXPHOS inhibition. Thus, targeting OXPHOS in Nrf2-mediated recurrent HPV-related HNSC in conjunction with cisplatin therapy may provide a synergistic precision target for improving oncologic outcomes in this devastating disease.

Recurrent HPV-related HNSC portends a poor prognosis (2). Previous research identified a predilection for alteration of oxidative stress genes (*KEAP1, NFE2L2*, or *CUL3*) among head and neck cancers of the classical subtype (7) and an association with poor survival in laryngeal SCC (21). The mesenchymal subtype has been associated with increased expression of innate immunity genes and a higher risk of nodal metastasis in oral cavity SCC (21). We found that 8/10 of the rOPSCCs were of the classical or mesenchymal subtype. In contrast, the majority of pOPSCCs were of the atypical or basal subtypes. The atypical subtype is associated with HPV-positive HNSC and activating mutations in exon 9 of the *PIK3CA* helical domain. The basal subtype is associated with *NOTCH1* inactivation and decreased *SOX2* expression.

Current treatment paradigms for recurrent head and neck cancer are limited and highly morbid. For resectable locoregional recurrence in the radiation therapy-naive patient, National Comprehensive Cancer Network (NCCN) guidelines (22) recommend surgery with adjuvant therapy depending on pathologic features versus multimodality chemoradiation. However, most patients (~95%) with HPV-related HNSC will already have received upfront radiation therapy administered as monotherapy or part of multimodality therapy (23). Balancing treatment morbidity with sound oncologic outcomes in previously irradiated patients is much more challenging. Treatment options include monotherapy or multimodality therapy with a combination of surgical extirpation, reirradiation, and/or systemic therapy versus palliative therapy or supportive care in some cases. Surgical management not infrequently entails morbid extirpation to ensure appropriate oncologic margins. Re-irradiation is associated with rates of grade ≥ 3 toxicity in close to 50% of patients (24) and increases the risk of adverse sequelae such as osteo- or soft-tissue necrosis. Thus, novel alternative therapeutic approaches should be explored.

Nrf2 is a biological double-edged sword acting both as a tumor suppressor and oncogene in part by regulating redox homeostasis which is a delicate balance. Nrf2 functions as a transcriptional regulator of antioxidant response element (ARE) genes and modulates mitochondrial function and OXPHOS efficiency abrogating the effect of ROS (25,26). Nrf2 affects the efficiency of OXPHOS (25) by raising the basal mitochondrial membrane potential, basal ATP levels, and oxygen consumption rates (27) while knockdown of Nrf2 decreases oxygen consumption and ATP production in cancer cells (28). Nrf2 provides substrate for OXPHOS and regulates the expression of complex IV cytochrome *c* oxidase subunits as well as nuclear respiratory factor 1 which regulates the expression of respiratory complexes (27,29-34). While protecting cells from oxidative stress, constitutive activation in multiple cancers promote a pro-survival phenotype through transcriptional induction of ROS-neutralizing and cytoprotective drug metabolizing enzymes (35).

Activation of the Nrf2 pathway induces expression of proteins involved in xenobiotic metabolism and clearance, inhibition of inflammation, repair and removal of damaged proteins, as well as transcription and activation of growth factors (36) – thus permitting cells to acquire features for therapeutic resistance. Upregulation of the Nrf2-mediated survival pathway protects tumor cells from chemotherapeutic agents including etoposide, doxorubicin and cisplatin (17,19,37,38). Constitutive activation of Nrf2 accelerates recurrence and induces metabolic reprogramming to re-establish redox homeostasis and upregulate *de novo* nucleotide synthesis in breast cancer cells (11). Ultimately this acquired Nrf2 activation confers lowered cancer therapeutic efficacy and materialization of therapeutic resistance. While much has been uncovered about the role of Nrf2 in therapeutic resistance, there is ample room for deepening our understanding of the mechanisms underlying this phenomenon.

Our *in vitro* and *in vivo* work describe for the first time, that Nrf2 drives the proliferation of HPV-driven cancers. Prior studies have found similar results. For example, Keap^-/-^ cells proliferate faster than parental cells while Nrf2^-/-^ cells proliferate more slowly (39,40) modulated by variation in growth factors (36). The Nrf2 gene contains a 12-O-tetradecanoylphorbol-13-acetate (TPA) response element (TRE) in its promoter (41) which is a binding site for the AP-1 transcription factor, an important modulator of cell proliferation where multiple oncogenic signals converge (42). Oncogenic proteins that regulate proliferation such as *KRAS, BRAF* and *MYC* increase the expression of Nrf2 (41,43), which corroborates our findings. In addition to regulating cell proliferation, Nrf2 plays a critical role in mitochondrial physiology and biogenesis in the context of other critical metabolic regulatory genes including Myc.

Mitochondrial respiration depends on Nrf2 activity. We found that complex I inhibitors IACS-010759 and rotenone decrease ATP production in Nrf2 overexpressing cells indicating a role for Nrf2 in mitochondrial respiration. These OXPHOS inhibitors also decreased cellular proliferation in a Nrf2-mediated manner indicating a Nrf2-dependent relationship between OXPHOS and the cell cycle. This suggests that in context of Nrf2 overexpression, glucose consumption and energy demand needs are met in part through activating OXPHOS. Abundance of Nrf2 also led to increased ROS production which may independently trigger cellular proliferation (44-46). Inhibition of mitochondrial respiration with NAC suggests that ROS is not only a byproduct of OXPHOS but also acts in a positive feedback mechanism to regulate OXPHOS.

In this study, we identified gene expression enrichment of Nrf2, OXPHOS, Myc, fatty acid metabolism, and DNA damage response signaling among metachronous recurrent HPV-related HNSCs. Prior work demonstrated that Myc activation reprograms cancer cell metabolism by activating genes involved with glycolysis, glutaminolysis, and mitochondrial biogenesis (47). Moreover, a substantial body of evidence illustrates the effect of Nrf2 in driving mitochondrial membrane potential, mitochondrial biogenesis, fatty acid oxidation, and OXPHOS (25,27,48,49). This led us to query existing datasets for a relationship between these pathways and overall survival. One group found an Nrf2-related gene expression signature associated with poor survival among the TCGA head and neck squamous cell carcinoma (HNSC) data and in multiple non-small cell lung cancer cohorts (50,51). Moreover, our analysis of the TCGA HNSC data re-capitulated and expanded upon these findings in that we observed a survival difference dependent on Nrf2 dysregulation at the genomic and pathway levels. In contrast, we did not identify a survival difference among the TCGA HPV-related HNSCs when stratified by Myc gene dysregulation, though at the Myc pathway level, there was an association with pathway enrichment and worse survival. This finding is not surprising in the context of the key role Myc plays as an oncogene including in the dysregulation of cell metabolism. OXPHOS pathway dysregulation was significantly associated with survival among the TCGA HPV-related HNSCs, though the group with the worst survival had an intermediate level of OXPHOS pathway enrichment. Lastly, GSEA of the Nrf2-upregulated TCGA HPV-related HNSCs demonstrated enrichment in OXPHOS signaling, glycolysis, fatty acid metabolism, *MYC, MTOR*, and ROS signaling pathways suggesting an interaction among these metabolic signaling pathways in Nrf2-altered tumors that ultimately converge on increase tumor cell fitness and worse overall patient survival.

To our knowledge, this is the first report identifying Nrf2-dependent OXPHOS inhibitor sensitization in head and neck cancer, a potential targetable lynchpin. Several studies have illustrated a role for either OXPHOS or Nrf2 in tumorigenesis. Nrf2 dysregulation in an esophageal cancer model demonstrated an association with increased cell proliferation and altered metabolism (52). Prior work analyzing the HPV-host protein network identified an HPV E1-KEAP1 interaction that phenocopies inactivating mutations in the KEAP1-Nrf2 pathway leading to the expression of cytoprotective genes (9). On the other hand, OXPHOS drives treatment resistance independent of Nrf2 as prior research in human ovarian cancer suggests that oxidative phosphorylation promotes platinum-based chemoresistance (53). Genomic analysis of mantle cell lymphoma found that metabolic reprogramming towards OXPHOS and glutaminolysis is associated with resistance to ibrutinib, and that resistance was overcome with OXPHOS inhibition via IACS-010759 (54). Complex I inhibition via IACS-010759 also delays regrowth of neoadjuvant chemotherapy-resistant triple-negative breast cancer (55). Lastly, complex I activity can signal antioxidant response through ROS-independent mechanisms using ERK5 as a signaling meditator (56). Taken together, there is much to be understood regarding the interaction of Nrf2 and OXPHOS in the overall metabolic signaling milieu and their role in treatment resistance. Our study illustrates Nrf2-dependent sensitization to OXPHOS inhibition.

Limitations of the current study include accounting for the impact of tumor heterogeneity on our expression results. Head and neck cancers harbor genomic heterogeneity accounting in part for their ability to resist our current treatment regimens (57). It is possible that other genomic, epigenomic, or virally-mediated (9) mechanisms are driving recurrence aside from those identified in our differential expression and gene set enrichment analyses and that we are capturing a subset of highly expressed genes in certain tumor clones. For example, the phosphoinositide 3-kinase and AKT signaling pathway is frequently enriched in HPV-related HNSCs and regulates redox metabolism in cancer (58,59). Evidence from the TCGA HNSC dataset demonstrate an association between Nrf2 dysregulation and survival lending support to the role of Nrf2 in recurrence. Additionally, other metabolic signaling cascades may be playing a role in driving recurrence warranting further analysis. There is also potential for misclassification of HPV-status given that we used p16 status as a surrogate marker for HPV-driven disease. Further work will aim to increase or understanding of mechanisms underlying Nrf2-dependent OXPHOS inhibitor sensitivity and evaluate the effects of metabolic adaptations on the tumor microenvironment apropos to driving treatment resistance.

In summary, we observed that metachronous recurrent HPV-related HNSCs are enriched in Nrf2 and OXPHOS signaling dysregulation in an overall metabolic dysregulated milieu and that Nrf2-enriched HPV-related HNSC displays increased sensitivity to OXPHOS inhibition. Treatment options for recurrent HPV-related HNSC are limited. IACS-010759 is currently under investigation in a phase I clinical trial for metastatic or unresectable malignancies (60). Our data may be used to support the rational use of OXPHOS inhibitors in patients with Nrf2-enriched, recurrent HPV-related HNSC cancer in a clinical trial providing expanded options for patients with this devastating disease.

## Methods

### Data collection

Clinical data from the University of Pittsburgh tumor samples were abstracted by study investigators.

Archival tissue specimens from the University of Pittsburgh tumor samples were processed via fixation in 10% neutral buffered formalin, dehydrated in ethanol and embedded with paraffin wax (FFPE). H&E slides were prepared, and areas with high tumor density (>75% tumor cells) were marked for extraction. Two-mm punch biopsies were taken from the FFPE tumor dense regions for downstream tumor RNA extraction. FFPE tissue was de-paraffinized with xylenes, washed in consecutive ethanol rinses (100% and 70%), and heated to remove formalin cross-linking (61).

Tumor RNA was extracted using the *Quick-RNA* FFPE Miniprep extraction kit (Zymo Research, Irvine, CA). RNA was quantified using the Qubit™ RNA High Sensitivity Assay Kit (Thermo Fisher Scientific, Rockford, IL). Sample integrity was evaluated using the Agilent Bioanalyzer RNA Nano and Pico kits (Agilent Technologies, Santa Clara, CA).

### RNA sequencing and alignment

Sequencing and alignment steps for TCGA data were described previously (7,62). TCGA RSEM normalized expression data were obtained through FireBrowse (http://firebrowse.org/; illuminahiseq_rnaseqv2-RSEM_genes_normalized). University of Pittsburgh sample libraries were manually prepared via standard protocols using the Illumina TruSeq RNA Exome Library Prep Kit (Illumina, San Diego, CA) and sequenced on an Illumina NextSeq sequencing system using NextSeq 500 Mid- and High-Output 150 cycle kits (75 bp, paired-end; Illumina, San Diego, CA). Quality control was performed on the raw reads using RNA-SeQC (v1.1.7). Of the original twelve sets of paired primary and metachronous recurrent tumors, gene expression data from two pairs were not included in the downstream analyses due to low quality. Pre-processed short reads were aligned to the human genome reference sequence assembly (GRCh37/hg19) with the STAR2 aligner using a two-pass procedure.

### RNA expression analyses

FeatureCounts from SubRead (v1.6.0) was used for counting reads. Differential expression analysis was subsequently performed using the Empirical Analysis of Digital Gene Expression Data in R program (edgeR) (63,64). Negative binomial generalized log-linear models were applied to the gene-wise read counts for the metachronous recurrent and primary tumors adjusting for baseline differences between patients. Likelihood ratio tests were performed comparing log_2_ counts per million (CPM) between metachronous recurrent and primary tumors. Differentially expressed genes, defined as |log_2_ (ratio)| ≥1 with the false discovery rate (FDR) set at 5%, were identified.

### Pathway and gene set enrichment analysis

Gene Set Enrichment Analysis (GSEA; v20.0.5) was performed using TCGA RSEM expression data and Pittsburgh CPM data in GenePattern with min and max gene set sizes set to 5 and 1500, respectively (65,66). Single sample GSEA version 10.0.3 (67) was implemented in GenePattern using log_2_-transformed, median-centered expression data with rank normalization. The PANTHER classification system was used for pathway analysis of the expression data (68-70).

### Molecular subtype analysis

Subtype analysis was performed using the centroids in the gene expression classifier presented by Walter et al (12) with a reduction from 838 genes to 696 genes in common with Pittsburgh data and the Walter et al centroids. Each tumor was then assigned to one of the four subtypes (basal, atypical, classical, mesenchymal) by identifying the nearest centroid using a correlation-based similarity approach.

### TCGA survival analysis

TCGA clinical and genomic data were downloaded from cBioPortal (71,72). Classification of samples by genomic aberration status (i.e., mutation, copy number variant, expression) was extracted from cBioPortal. For classification by gene set cluster, TCGA RSEM normalized expression data were clustered using the Consensus Cluster Plus algorithm to facilitate gene expression cluster assignment (73). Kaplan-Meier survival functions were stratified either by genomic aberration status or expression cluster status and estimated using the Survminer package in R (74). In order to manage confounding, Cox proportional hazards models were fit controlling for age and tumor stage (75) and forest plots used to present the hazard ratios.

### Transcription factor/motif activity analysis

To analyze activities of transcription factor binding motifs (TFBM) using RNA-seq data, we used the Integrated System for Motif Activity Response Analysis (ISMARA) (13).

### Cell lines and tumor samples

HPV-16 expressing HNSC cell lines used in the present study were: UMSCC47 (isolated from the primary tumor of the lateral tongue of a male patient, established by Dr. Thomas Carey, University of Michigan), UPCI-SCC90 (from tongue squamous cell carcinoma (isolated by Dr. Susanne Gollin, University of Pittsburgh) and 93VU147T (squamous cell carcinoma isolated from the floor of mouth). UMSCC47 and 93VU147T cells were maintained in high glucose Dulbecco’s Modified Eagle’s Medium (DMEM) while UPCI-SCC90 was cultured in Eagle’s Minimum Essential Medium (EMEM) supplemented with 10%FBS and 1% penicillin/streptomycin mixture. Cells were used for collecting data for 10 passages after thawing and then discarded. All cell lines were authenticated using human cancer cell line STR profiles. All cell lines were maintained at 37°C in a humidified atmosphere of 95% air and 5% CO2. All human tumor samples were obtained from the University of Pittsburgh Medical Center in accordance with established University of Pittsburgh IRB guidelines.

### Reagents and Chemicals

DMEM and EMEM were purchased from GE Healthcare Life Sciences (Logan, UT). Fetal Bovine Serum (FBS) was obtained from Gemini Bioproducts (West Sacramento, CA). Penicillin/streptomycin antibiotic mixture was purchased from Invitrogen-Life Technologies (now part of Thermo Fisher Scientific, Waltham, MA). Dimethyl sulfoxide (DMSO), *N*-acetyl-L-cysteine (NAC), trigonelline, and rotenone were from Sigma-Aldrich (St. Louis, MO). Cisplatin was from EMD Millipore (now part of Millipore Sigma, Burlington, MA). Apocynin was purchased from Santa Cruz Biotechnology (Dallas, TX). IACS-010759 was purchased from Selleckchem (Houston, TX).

Stock solution of each compound was stored at -20°C and diluted in culture media before use. Antibody against Nrf2, HMOX-1, NQO1 and β-tubulin were from Abcam (Cambridge, MA). Primers were bought from IDT Technologies (Coralville, IA). Nrf2 endoribonuclease-prepared small interfering RNA (esiRNA) was purchased from Sigma-Aldrich (St. Louis, MO). Kits for RNA isolation and QIAshredder were purchased from Qiagen (Valencia, CA). iScript RT supermix for RT-qPCR and iQ™ SYBR^®^ Green Supermix for qPCR was from Bio-Rad Laboratories (Hercules, CA).

### Nrf2 overexpressing cell lines

The Nrf2-expressing retrovirus (MSCV-Nrf2) was a kind gift from Dr. David Tuveson’s laboratory (Cold Spring Harbor, NY). UMSCC47 and UPCI-SCC90 cells were stably infected with retrovirus expressing empty vector (MSCV-Neo) or Nrf2 (MSCV-Nrf2). Cells were selected in culture with media supplemented with 200 μg G418 for more than 4 weeks. Nrf2 overexpression was confirmed by qPCR.

### Nrf2 esiRNA transfection

Cells were plated in 6-well plates and grown to 70% confluency. Ten μM siRNA was transfected with Lipofectamine RNAiMAX reagent (Invitrogen) according to manufacturer’s instructions before re-plating and analyzing for qPCR or cell proliferation.

### Colony formation assay

About 2×10^3^ cells were plated in 12-well plates and left in the incubator for 10-12 days for the development of colonies with or without indicated treatments. Media was changed twice weekly and cultures were monitored for 7-14 days (depending on growth rate differences) to allow for Neo cells to reach >50 cells per colony. They were fixed in 4% buffered formalin and stained with crystal violet. The plates were scanned and analyzed in Image J using the colony area plugin.

### Western blotting

Cells were collected and lysed using appropriate amount of Tris–HCl/EDTA buffer supplemented with protease and phosphatase inhibitor. Lysates were incubated on ice for 15-20 minutes, sonicated and spun down at maximum speed for 20 mins. After protein quantification (Bradford’s method), lysates were prepared in β-ME and boiled. Between 50-80 μg protein was loaded in each lane. Tubulin or vinculin normalization was performed for each experiment. Immunoreactive bands were visualized and quantified using the LiCor Odyssey system.

### Quantitative real-time PCR (qRT-PCR)

Total RNA from 93VU147T or Nrf2 overexpressing UMSCC47 and UPCI-SCC90 cells were isolated using RNeasy kit. Tumors were disrupted and homogenized in the appropriate volume of lysis buffer provided in the RNeasy kit. The tissue lysate was loaded onto the QIAshredder homogenizer after which RNA was isolated using the manufacturer’s protocol. First-strand cDNA was synthesized using iScript RT and qPCR was done using suitable dilution of cDNA. RT conditions: 15 sec denaturation/95°C, 30 sec annealing/60°C, and 30 sec extension/72°C for 40 cycles. Relative quantification was performed using the 2^−ΔΔCq^ method.

Primer sequences for target genes are as follows:

Nrf2
Forward 5’-CGG TAT GCA ACA GGA CAT TG-3’
Reverse 5’-ACT GGT TGG GGT CTT CTG TG-3’
HMOX1
Forward 5’-GAGACGGCTTCAAGCTGGTGAT-3’
Reverse 5’-CCGTACCAGAAGGCCAGGTC-3’
NQO1
Forward 5’-CGC AGA CCT TGT GAT ATT CCA G-3’
Reverse 5’-CGT TTC TTC CAT CCT TCC AGG-3’

### Cell proliferation assay

We plated 2.5 – 5×10^4^ cells in 96-well plates and allowed to attach overnight. After indicated treatments, 10 μl premix WST-1 cell proliferation reagent (Takara Bio Inc, Clontech Laboratories, Inc.) was added to each well, and the plate was returned to incubator for 2 hours after which absorbance was read at 450 nm in a microplate reader (BioTek Instruments Inc).

### Amplex Red assay

UMSCC47 and UPCI-SCC90 cells expressing Neo and Nrf2 cells were used to determine the extracellular H_2_O_2_ using 10-acetyl-3,7-dihydroxyphenoxazine (Amplex Red Hydrogen Peroxide Assay kit from Molecular Probes Inc., Eugene, OR) following the manufacturer’s instructions. Approximately 25×10^3^ cells/well were plated in 96-black well plates and allowed to attach overnight. After NAC (20 mM) or apocynin (100 μM) treatment for 2-3h, cells were washed once in PBS, then incubated for 30 minutes at 37°C in reaction buffer containing 0.2 U/mL horseradish peroxidase and 100 μM Amplex Red. Wells containing all reactants except samples were run in parallel to account for background fluorescence. Fluorescence was expressed as H_2_O_2_ released using a standard curve generated with known concentrations of H_2_O_2_ stabilized solution using the Synergy H1 Hybrid microplate reader from BioTek at excitation of 530 nm and emission of 590 nm. Cells were then counterstained with crystal violet to correct the H2O2 released values for variations in cell culture densities.

### MitoSOXassay

Detection of mitochondrial superoxide in live cells was done using MitoSOX™ Red (Molecular Probes Inc., Eugene, OR). We cultured 25,000 cells in 96-well plate and washed the cells with PBS, then incubated the plates with 5 μM MitoSOX in PBS for 30 minutes at 37°C. Following incubation in the dark, MitoSOX was removed, cells were washed again with PBS and then plates were read in Synergy H1 Hybrid microplate reader from BioTek at excitation of 510 nm and emission of 595 nm.

### ATP production

The ATP-monitoring ATPlite luminescence assay system from Perkin Elmer Inc. (Waltham, MA) was used for quantitative evaluation of proliferation in cultured UMSCC47 and UPCI-SCC90 cells. Briefly, 25×10^3^ cells/well were plated in 96-black well plates and allowed to attach overnight. After appropriate treatments, luminescence was read on a Synergy H1 Hybrid microplate reader (BioTek). ATP production was measured by ATPlite™ assay using standard curve of ATP and further normalization with cell count. IACS-010759 applied at 10nM (24h), rotenone at 10μM (2h followed by rescue in regular DMEM media for 24h) and 20μM FCCP (30’). Each experiment was repeated three times and combined data of mean ± SEM are illustrated.

### Tetramethylrhodamine ethyl ester (TMRE) assay

The TMRE-Mitochondrial Membrane Potential Assay Kit (Abcam, Cambridge, MA) was used for quantifying changes in mitochondrial membrane potential in live cells using a Synergy H1 Hybrid microplate reader (BioTek, Winooski, VT). The assay implements FCCP (carbonyl cyanide 4-(trifluoromethoxy) phenylhydrazone), an ionophore uncoupler of oxidative phosphorylation, as positive control. Adherent and live cells are stained for 15-20 minutes with or without FCCP, after which TMRE staining is measured by microplate spectrophotometry (excitation/emission 549/575 nm).

### In vivo xenograft studies

The animal experiment protocol was approved by the Institutional Animal Care and Use Committee of the University of Pittsburgh. A total of 2×10^6^ UMSCC47-Neo and UMSCC47-Nrf2 overexpressing cells mixed with matrigel were injected subcutaneously into either flank of NOD SCID mice aged 5-8 weeks (N=6-8 tumors/ group). Tumor volume was measured once a week using calipers and was calculated using the formula: 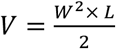 (76). Two-way ANOVA with Sidak’s test used to test for differences in mean tumor volume between Neo and Nrf2 groups over time. Bars represent the mean ± SEM. *, p-value < 0.01; ***, p-value < 0.0001.

### Statistics

R programming software (version 3.6.1) was used to perform statistical analyses (77). To account for multiple hypothesis testing, the false discovery rate was controlled using the method of Benjamini and Hochberg. In GSEA analyses, hypothesis generating q-values < 0.25 were used to determine if findings were statistically significant. In survival analyses, log-rank test p-values less than 0.05 were used to determine if findings were statistically significant. All other analyses used an alpha of 0.05 to test for statistical significance. All *in vitro* experiments were repeated 2-3 times. Representative data from CFA are shown with quantitation from all experiments. Difference between group means was tested with Student’s t-test. *, p-value < 0.05; **, p-value < 0.001; ***, p-value < 0.0001.

### Study approval

This study protocol was reviewed and approved by the University of Pittsburgh Institutional Review Board (IRB 99-069). Written consent was obtained for genomic characterization of tumor tissues for all participants prior to inclusion in this study. This study abided by the Declaration of Helsinki principles.

## Author Contributions

U.D., E.M., R.A.H., D.F., and A.V. conceived the experiment. M.K., D.F., and A.V. collected the data. R.A.H., A.V., Q.Z., D.F., and D.P. analyzed the data. R.A.H., A.V., Q.Z., D.F., D.P., H.O., and U.D. interpreted the data. R.A.H. and A.V. drafted the article. All authors participated in critical revision of the article. R.A.H., A.V., and U.D. were involved in final approval of the article.

## Acknowledgements

We thank Dr. Jeffrey Delrow of the Fred Hutchinson Cancer Research Center for bioinformatics analysis design support. We thank Dr. William LaFramboise for RNA sequencing support.

