## Supplemental Figures for "Recurrent human papillomavirus-related head and neck cancer undergoes metabolic re-programming and is driven by oxidative phosphorylation"

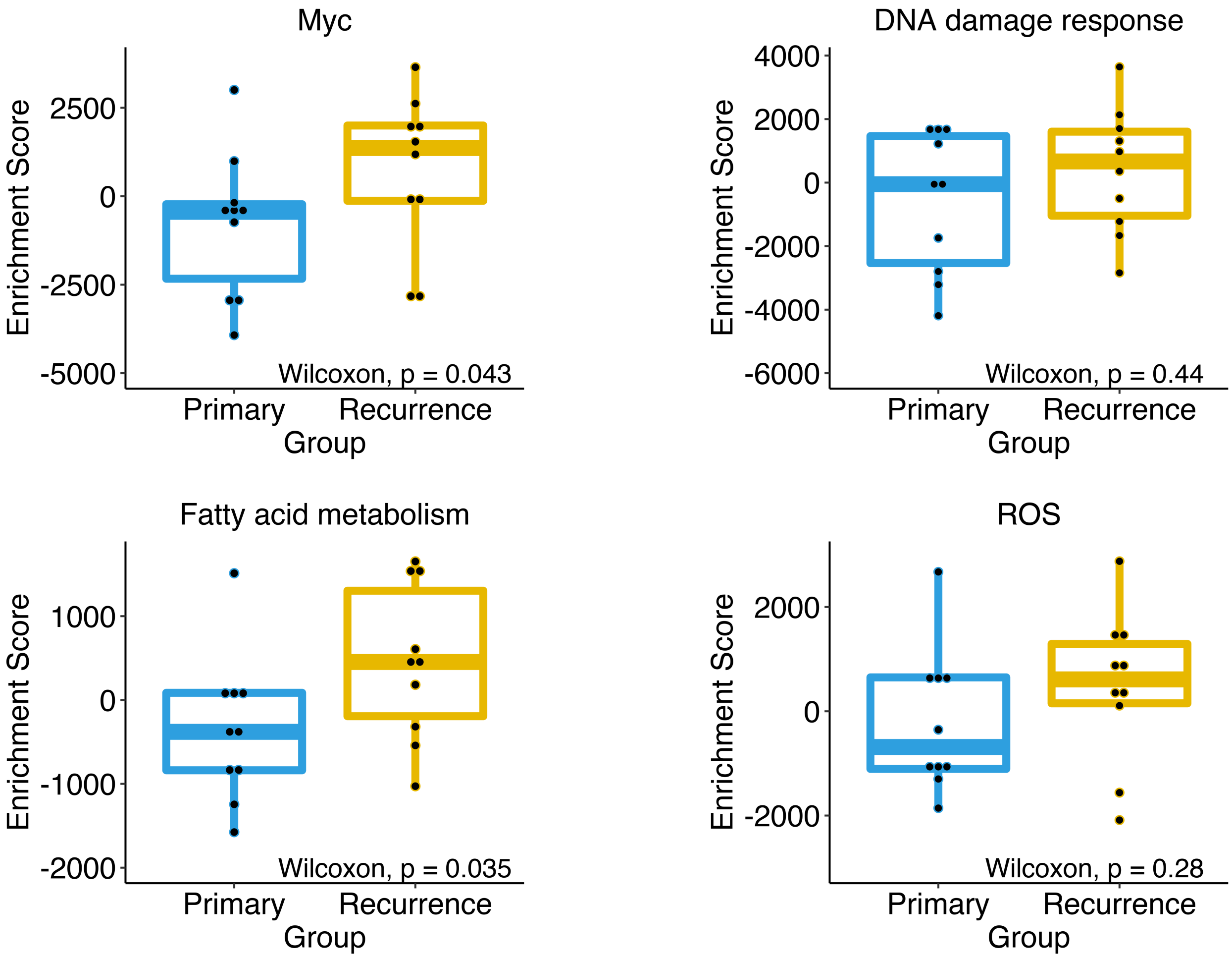


**Supplemental Figure 1. Single sample gene set enrichment analysis (ssGSEA) scores comparing primary (pOPSCC) versus metachronous recurrent (rOPSCC) HPV-related OPSCC.** Boxplots represent the median, $25^{th}$ and $75^{th}$ percentiles. The whiskers represent 1.5 * the interquartile ratio. Data represent the enrichment score (ES) for each sample (pOPSCC n=10; rOPSCC n=10). *Myc*, hallmark *MYC* v1 gene set. *DNA damage response*, Reactome p53-dependent DNA damage repair gene set. *Fatty acid metabolism*, hallmark fatty acid metabolism. *ROS*, hallmark reactive oxygen species gene set.

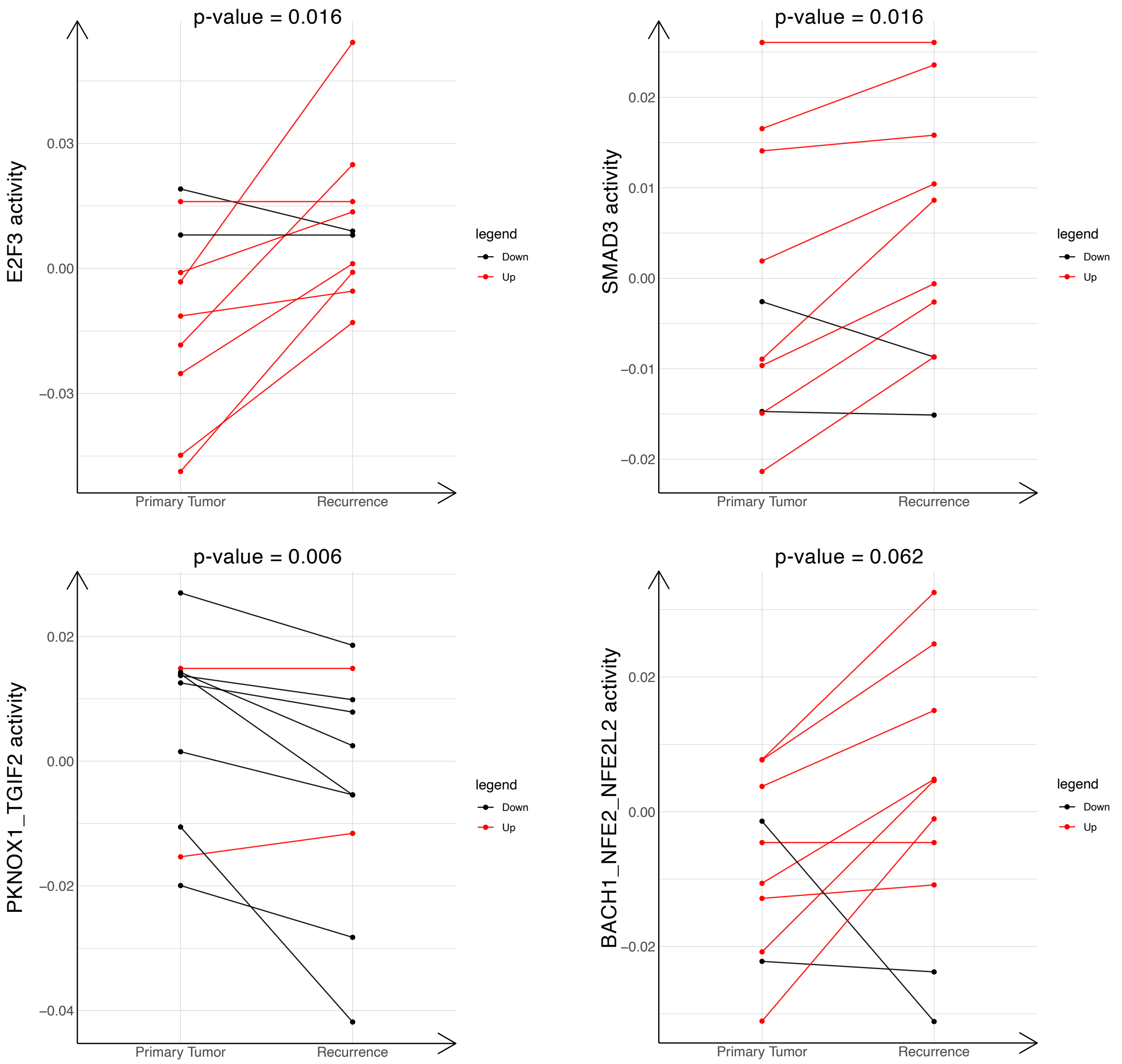


**Supplemental Figure 2. Inferred transcription factor/motif activity analysis of HPV-related pOPSCC vs rOPSCC**. Plots represent transcription factor activity in the primary and recurrent tumor samples. Transcription factor activity that increased in the rOPSCC compared to the pOPSCC denoted in red. Transcription factor activity that decreased in the rOPSCC compared to the pOPSCC denoted in black. One-sided Wilcoxon test p-values denoted at top of plot comparing pOPSCC vs rOPSCC.

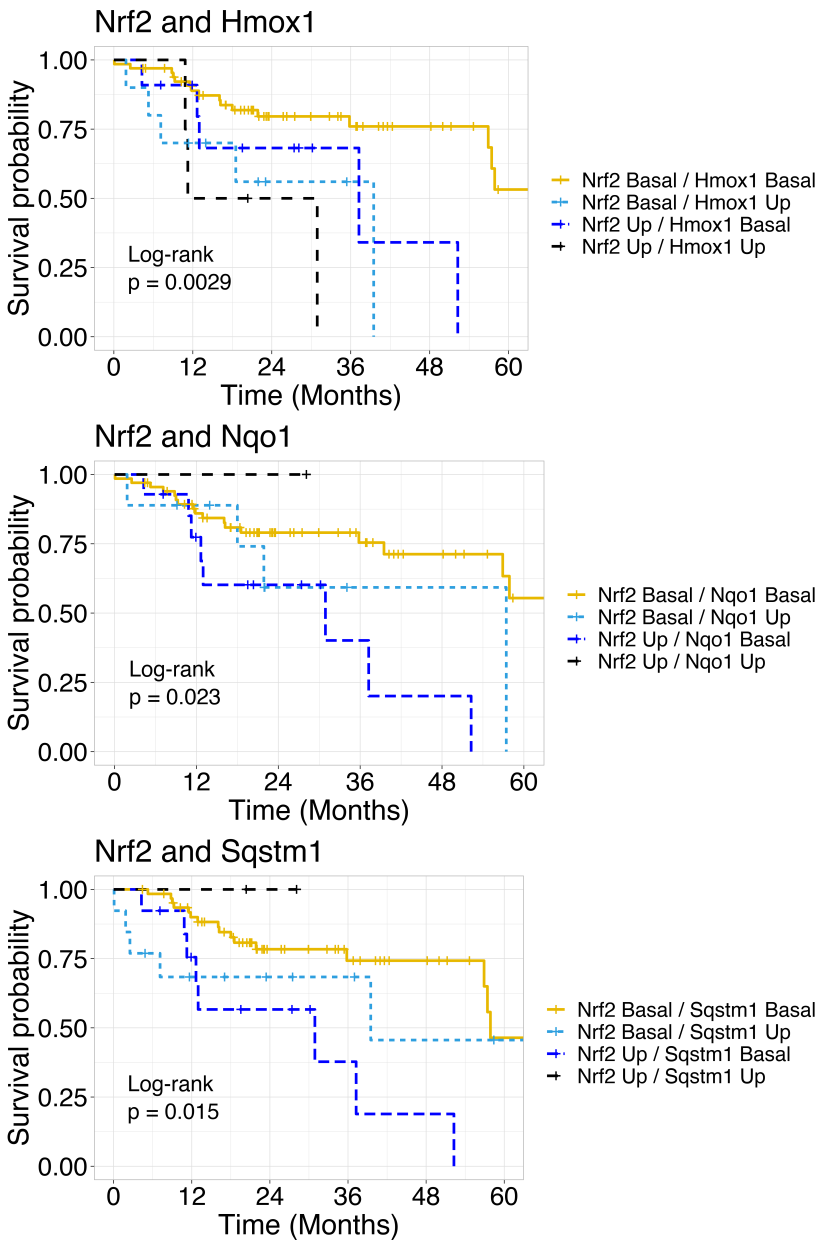


**Supplemental Figure 3. Survival among TCGA HPV-related OPSCC based on Nrf2 genomic alteration status in combination with genomic alterations of Nrf2 target gene alterations.** *Top panel*: Survival stratified by Nrf2 ± Hmox1 genomic alteration status. *Middle panel*: Survival stratified by Nrf2 ± Nqo1 genomic alteration status. *Lower panel*: Survival stratified by Nrf2 ± Sqstm1 genomic alteration status. Nrf2 Up: gene expression > 2, mutation, copy number gain or amplification; Nrf2 Basal: gene expression ≤ 2, no mutation, copy number neutral. Hmox1/Nqo1/Sqstm1 Up: gene expression > 2, copy number gain or amplification; Hmox1/Nqo1/Sqstm1 Basal: gene expression ≤ 2, copy number neutral.

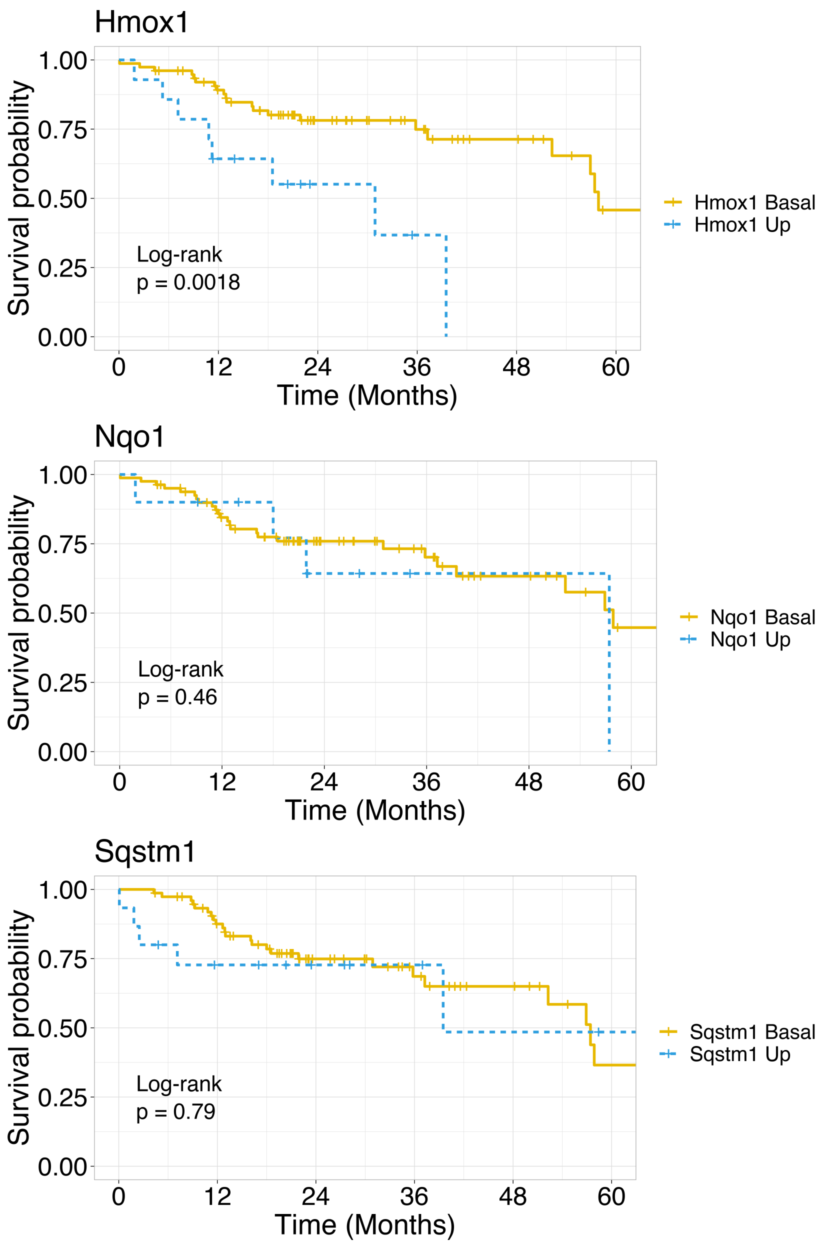


**Supplemental Figure 4. Survival among TCGA HPV-related OPSCC based on genomic alteration of Nrf2 target genes.** *Top panel*: Survival stratified by Hmox1 genomic alteration status. *Middle panel*: Survival stratified by Nqo1 genomic alteration status. *Lower panel*: Survival stratified by Sqstm1 genomic alteration status. Hmox1/Nqo1/Sqstm1 Up: gene expression > 2, copy number gain or amplification; Hmox1/Nqo1/Sqstm1 Basal: gene expression ≤ 2, copy number neutral. Log-rank global p-values labeled in plots.

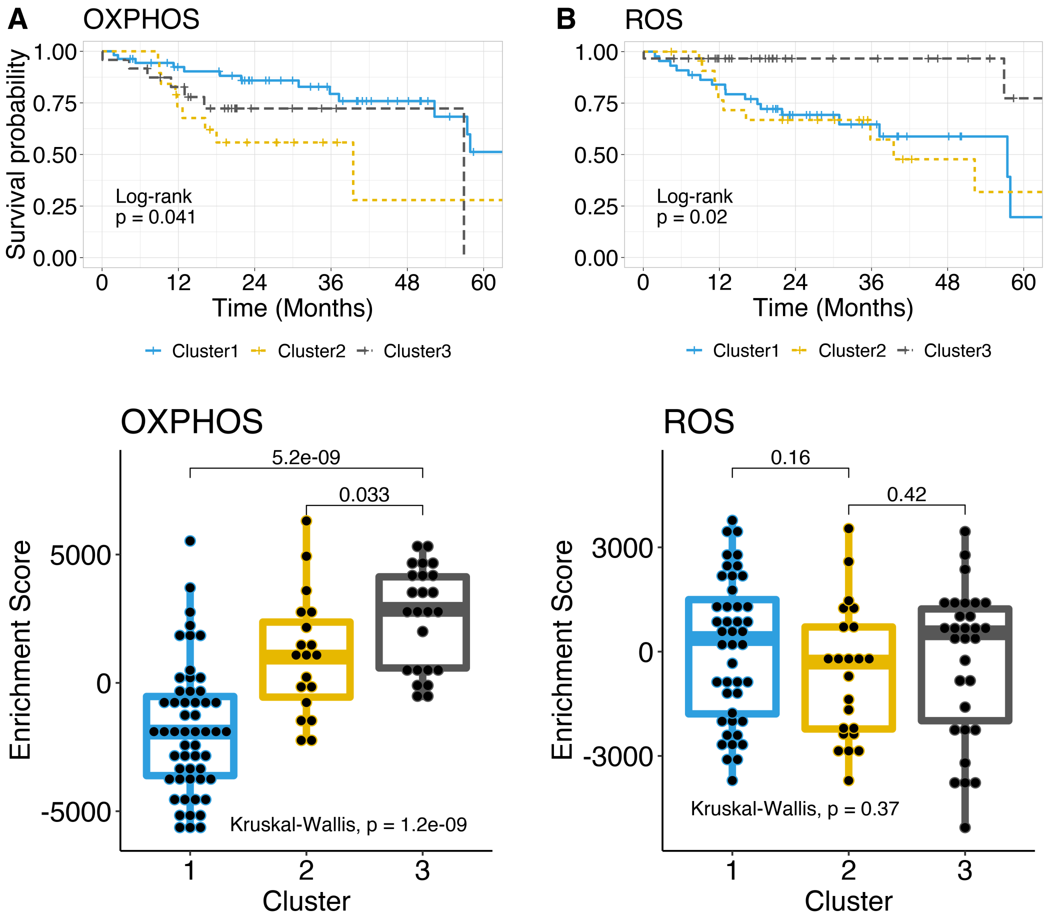


**Supplemental Figure 5. Survival analysis stratified by OXPHOS or ROS gene set expression clustering among TCGA HPV-related HNSC.** (**A**) *Upper panel*: Survival stratified by hallmark OXPHOS gene set expression cluster analysis. *Lower panel*: ssGSEA enrichment scores stratified by gene set expression cluster membership as assigned in clustering analysis. (**B**) *Upper panel*: Survival stratified by Hallmark reactive oxygen species (ROS) gene set expression cluster analysis. *Lower panel*: ssGSEA enrichment scores stratified by gene set expression cluster membership. Log-rank p-values labeled on each survival plot. Kruskal-Wallis test utilized for comparing enrichment scores between clusters. Values above the brackets between groups represent Wilcoxon rank-sum p-values.

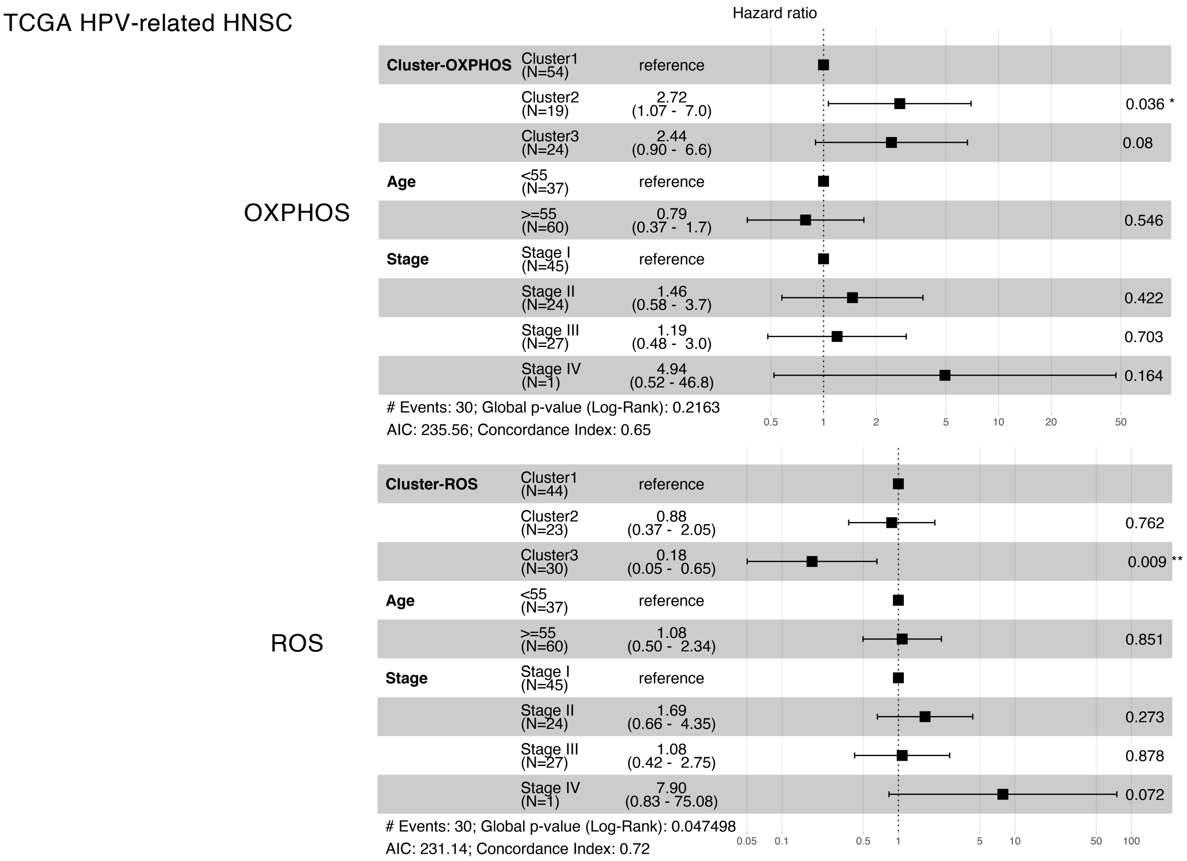


**Supplemental Figure 6. Forest plots comparing hazard ratios between groups stratified by OXPHOS or ROS gene set expression cluster, age, and tumor stage among TCGA HPV-related HNSC.** Cox regression hazard ratios by gene set expression cluster, age group, and tumor stage (AJCC 8th edition, clinical staging). Black boxes represent the hazard ratio with 95% confidence intervals represented by whiskers. P-values for each hazard ratio by predictor are represented on the far right of the plot. *, p-value < 0.05; **, p-value < 0.01. *OXPHOS*, hallmark oxidative phosphorylation; *ROS*, hallmark reactive oxygen species.

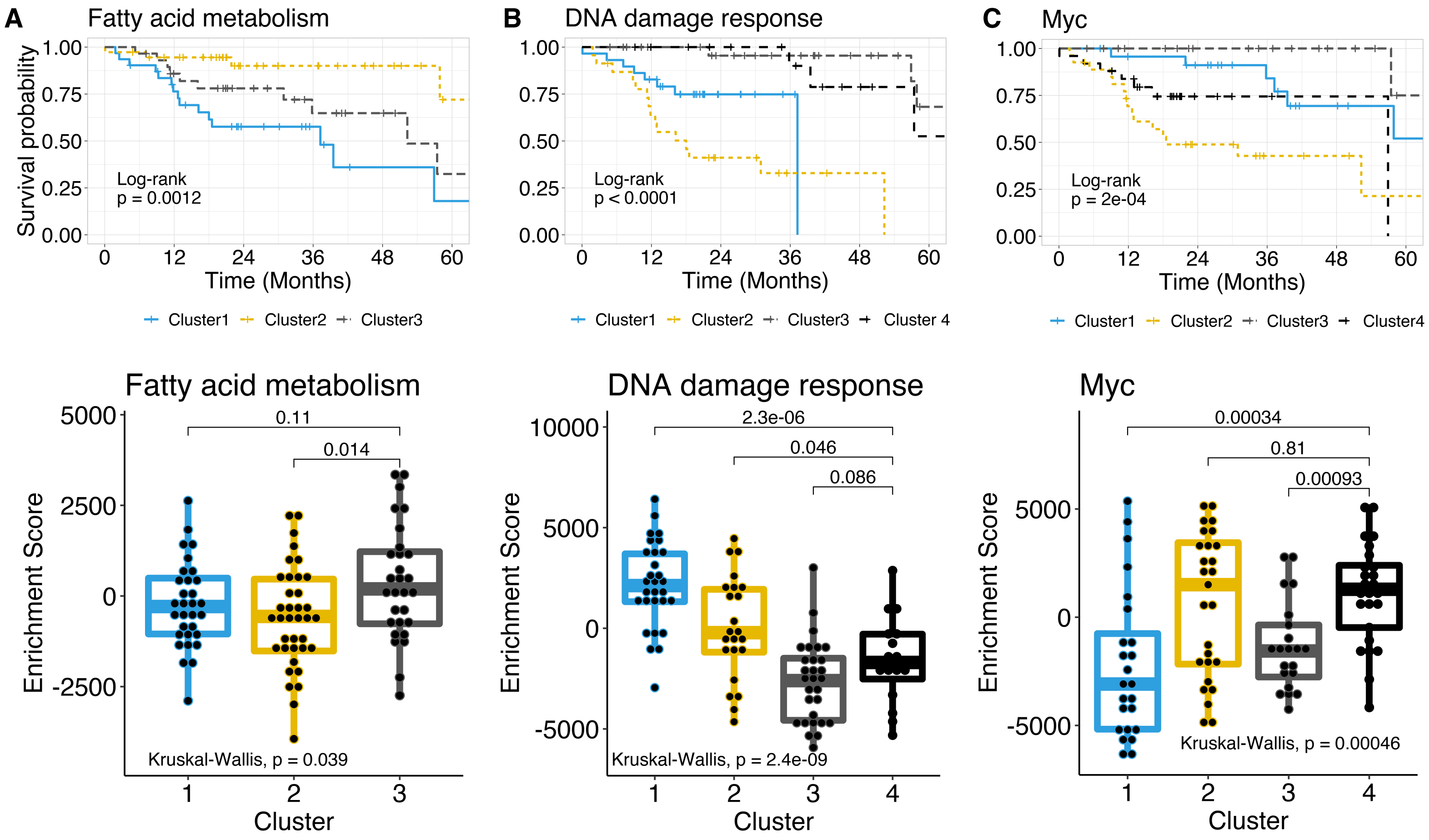


**Supplemental Figure 7. Survival analysis stratified fatty acid metabolism, DNA damage response, or Myc gene set clustering among TCGA HPV-related HNSC.** (**A**) *Upper panel*: Survival stratified by hallmark fatty acid metabolism gene set expression cluster analysis. *Lower panel*: ssGSEA enrichment scores stratified by gene set expression cluster membership. (**B**) *Upper panel*: Survival stratified by Reactome p53-dependent DNA damage repair gene set expression cluster analysis. *Lower panel*: ssGSEA enrichment scores stratified by gene set expression cluster membership. (**C**) *Upper panel*: Survival stratified by hallmark *MYC* v1 (Myc) gene set expression cluster analysis. *Lower panel*: ssGSEA enrichment scores stratified by gene set expression cluster membership. Log-rank p-values labeled on each survival plot. Kruskal-Wallis test utilized for comparing enrichment scores between clusters. Values above the brackets between groups represent Wilcoxon rank-sum p-values.

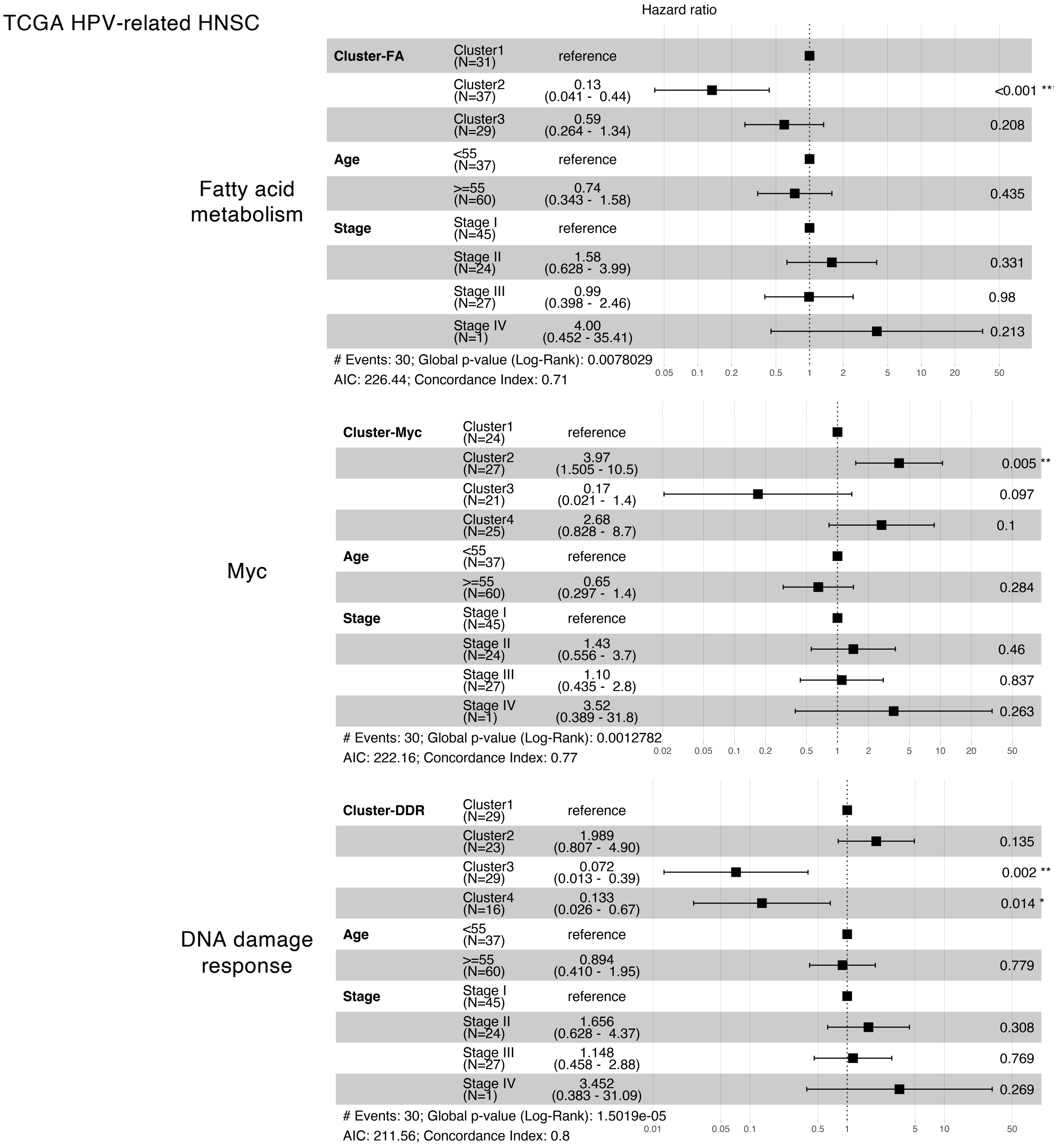


**Supplemental Figure 8. Forest plots comparing hazard ratios between groups stratified by fatty acid metabolism, Myc, or DNA damage response gene set expression cluster, age, and tumor stage among TCGA HPV-related HNSC.** Cox regression hazard ratios by gene set expression cluster, age group, and tumor stage (AJCC 8$^{th}$ edition, clinical staging). Black boxes represent the hazard ratio with 95% confidence intervals represented by whiskers. P-values for each hazard ratio by predictor are represented on the far right of the plot. *, p-value < 0.05; **, p-value < 0.01; ***, p-value < 0.001. *Fatty acid metabolism*, hallmark fatty acid metabolism; *Myc*, hallmark *MYC* v1; *DNA damage response*, Reactome p53-dependent DNA damage repair.

A

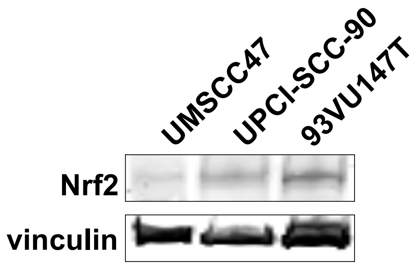


B

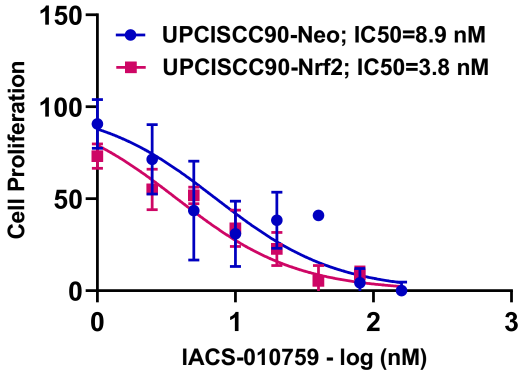


C

| Cell type | IC50 (nM) of IACS |
|---|---|
| UMSCC47-Neo | 8.9 |
| UMSCC47-Nrf2 | 1.6 |
| UPCI-SCC90-Neo | 8.9 |
| UPCI-SCC90-Nrf2 | 3.8 |
| UPCI-UDSCC17-70 | 2.8 |

D

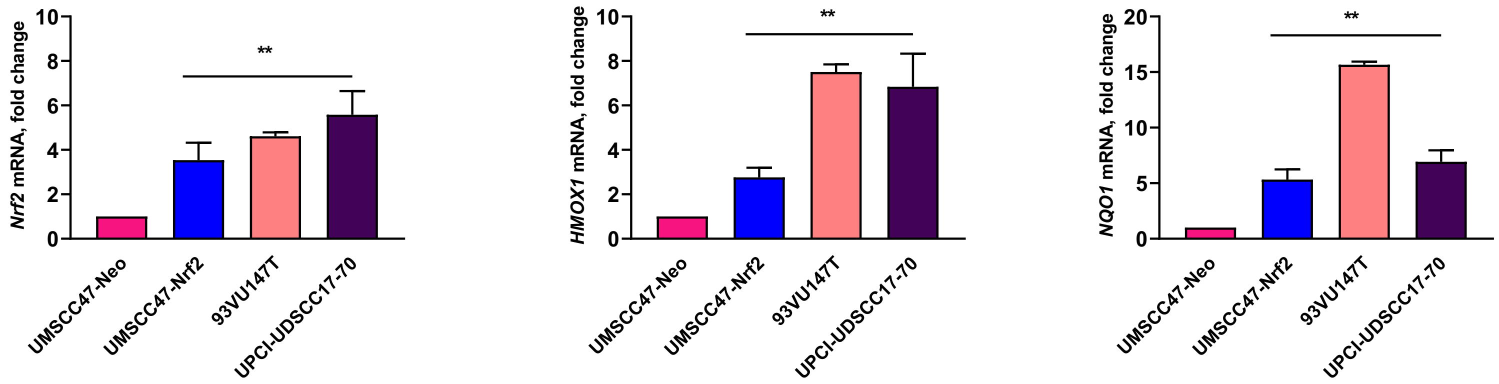


**Supplemental Figure 9. Mitochondrial complex I inhibitor IACS-010759 sensitizes Nrf2-overexpressing HPV-positive HNSC cells and Nrf2 activation pathway in panel of cells** (**A**) Western blot for Nrf2 in parental cell lines. (**B**) Cell proliferation was measured using WST-1 reagent in UPCISCC90-Neo and -Nrf2 cells treated with increasing concentrations of IACS-010759 for 24h. Experiments were performed three times and representative data from one experiment is shown. (**C**) Table showing differential IC50 for IACS-010759 between cell lines. (**D**) qPCR for *Nrf2*, *HMOX1* and *NQO1* in panel of cell lines. Experiments were repeated two or three times and combined data is shown as mean ± SEM. Differences are calculated using one-way ANOVA with Dunnet's test. **p- value < 0.01.

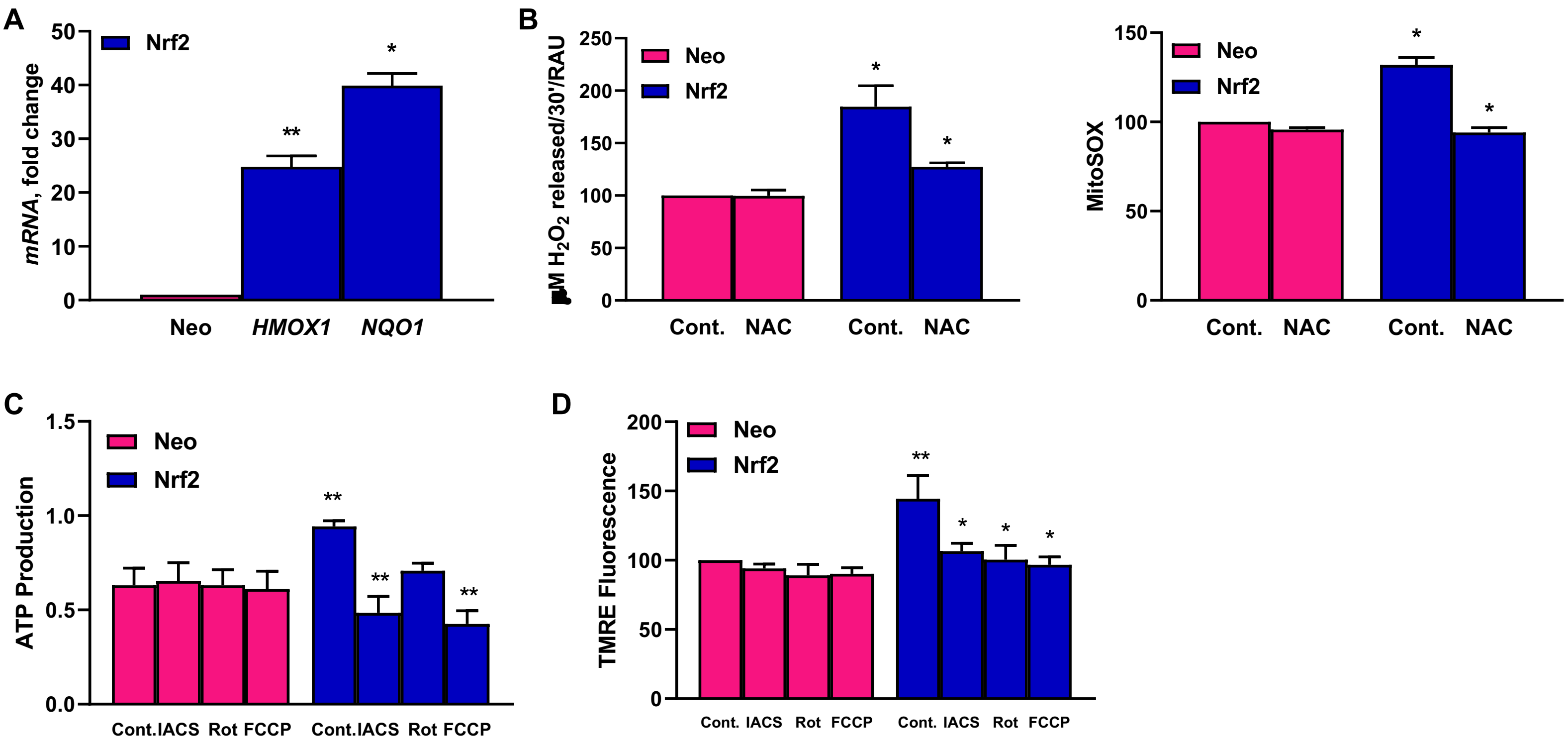


**Supplemental Figure 10. Nrf2 promotes cell proliferation via OXPHOS in HPV-positive UPCISCC90-Nrf2 cells.** (**A**) qPCR for *HMOX1* and *NQO1*. Student's t-test used to compare fold change relative to Neo. **, p-value < 0.001; ***, p-value < 0.0001. (**B**) *Left panel*: ROS measured using Amplex Red in cells treated with and without 10mM NAC for 3h as $H_2O_2$ released in 30'. *Right panel*: Mitochondrial superoxide is measured in control and in 10mM NAC (3h) treated cells using MitoSOX™ Red reagent. Differences relative to Neo control calculated using one-way ANOVA with Dunnet's test. *, p-value < 0.05. (**C**) ATP production measured using ATPlite™ using standard curve of ATP and further normalization with cell count. (**D**) Mitochondrial membrane potential measured using TMRE dye. IACS-010759 used at 10nM (24h), rotenone at 10µM (2h followed by rescue in regular DMEM media for 24h and 20µM FCCP for 30'. Treatment groups in Nrf2 cells are compared to Nrf2 control, and the Nrf2 control group is compared to Neo control. Differences are computed by one-way ANOVA with Tukey's test. **, p-value < 0.05; *, p-value < 0.005. Results represent mean ± SEM. Experiments repeated three times and combined graph of n=3 is shown.
